# Dissecting Multiparametric Cerebral Hemodynamics using Integrated Ultrafast Ultrasound and Multispectral Photoacoustic Imaging

**DOI:** 10.1101/2023.11.07.566048

**Authors:** Haoyang Chen, Shubham Mirg, Prameth Gaddale, Sumit Agrawal, Menghan Li, Van Nguyen, Tianbao Xu, Qiong Li, Jinyun Liu, Wenyu Tu, Xiao Liu, Patrick J. Drew, Nanyin Zhang, Bruce J. Gluckman, Sri-Rajasekhar Kothapalli

## Abstract

Understanding brain-wide hemodynamic responses to different stimuli at high spatiotemporal resolutions can help study neuro-disorders and brain functions. However, the existing brain imaging technologies have limited resolution, sensitivity, imaging depth and provide information about only one or two hemodynamic parameters. To address this, we propose a multimodal functional ultrasound and photoacoustic (fUSPA) imaging platform, which integrates ultrafast ultrasound and multispectral photoacoustic imaging methods in a compact head-mountable device, to quantitatively map cerebral blood volume (CBV), cerebral blood flow (CBF), oxygen saturation (SO2) dynamics as well as contrast agent enhanced brain imaging with high spatiotemporal resolutions. After systematic characterization, the fUSPA system was applied to quantitatively study the changes in brain hemodynamics and vascular reactivity at single vessel resolution in response to hypercapnia stimulation. Our results show an overall increase in brain-wide CBV, CBF, and SO2, but regional differences in singular cortical veins and arteries and a reproducible anti-correlation pattern between venous and cortical hemodynamics, demonstrating the capabilities of the fUSPA system for providing multiparametric cerebrovascular information at high-resolution and sensitivity, that can bring insights into the complex mechanisms of neurodiseases.

## 1 Introduction

Cerebral hemodynamics, controlled by various physiological factors like autoregulation, neurovascular coupling, and vascular reactivity, is vital for brain health as well as the exchange of gases and nutrients [1, 2]. Cerebral blood volume (CBV), cerebral blood flow (CBF), and cerebral oxygen saturation (SO2) are three important hemodynamic parameters commonly studied for assessing cognitive function and help gain insights into the diagnosis and treatment of neurological disorders [3, 4]. However, current modalities fail to provide these individual cerebral hemodynamic information in the deep brain. Optical imaging methods, such as two-photon fluorescence microscopy and laser speckle contrast imaging [5, 6], can map changes in CBF and individual vessel diameter in the rodent brain with high resolution. Yet the imaging depth is limited to superficial regions (< 2 mm inside the brain) and often requires surgical procedures to create optically transparent windows [7, 8]. On the other hand, functional magnetic resonance imaging (fMRI) based blood oxygen level-dependent (BOLD) signal is widely used in neuroscience to non-invasively study whole brain hemodynamic changes [9]. However, fMRI exhibits low spatial resolution and the BOLD signal is a mix of several hemodynamic factors [9, 10], making interpretation confounding. Moreover, the noise and the bulky size of fMRI instrumentation restrict its applicability in studies involving naturally behaving animals [10].

The last decade has seen rapid advances in multispectral photoacoustic (MSPA) and ultrafast ultrasound (UFUS) imaging techniques that overcome the above discussed limitations of optical imaging and fMRI [11–13]. MSPA is a hybrid imaging technique in which optical pulses of different wavelengths illuminate tissue of interest, subsequently these light pulses are converted to broadband ultrasound waves by light absorbing molecules such as hemoglobin. Based on the differential optical absorption of oxy-hemoglobin (HbO) and deoxy-hemoglobin (HbD), MSPA can map time-resolved total hemoglobin concentration and oxygen saturation (SO2) changes at ultrasound resolutions [14–16]. In addition to the label-free vascular information, MSPA can provide a wide range of exogenous molecular contrast via the administration of optical contrast agents such as indocyanine green (ICG) [17] or genetically encoded light absorbing proteins [18]. In contrast, UFUS imaging provides microvasculature information based on ultrasound scattering from moving particles such as red blood cells or injected microbubbles. Power Doppler ultrasound imaging (also known as functional ultra-sound - fUS) uses UFUS based multiangle coherent plane wave compounding [11, 19, 20] to map changes in CBV with high spatial (100 µm × 100 µm × 300 µm voxel) and temporal (300 msec – 1 sec) resolutions, and has shown good agreement with direct measures of hemodynamics at the single vessel level using 2-photon microscopy [21, 22]. On the other hand, by fitting autocorrelation functions to the speckles of UFUS compounded frames, a quantitative assessment of CBF change can be obtained [23]. To help delineate different vascular structures, the functional CBF information can be further differentiated into ascending flows and descending flows based on the extracted positive and negative frequencies. In addition, by intravenously injecting acoustic contrast agents (microbubbles - MBs) and applying MB localization and tracking methodologies, UFUS provides super-resolved ultrasound localization microscopy (ULM) images of vascular anatomy, vertical blood flow, and flow speed with a spatial resolution that is beyond the ultrasound wave diffraction limit [12, 24]. However, despite these techniques showing promise for hemodynamic imaging, each technique in isolation is limited in the scope of information it can provide as they are not obtained concurrently. Thereby necessitating the development of a multimodal imaging setup for understanding the individual while interdependent hemodynamic parameters [2], especially in an organ as complex as the brain.

Towards this, recent studies have reported combined ultrasound and photoacoustic technologies for high-resolution functional brain imaging [25–27]. However, no studies integrated the MSPA and UFUS approaches for comprehensive analysis of individual hemodynamic parameters (CBV, CBF, and SO2) of deep brain in response to different stimuli. Here, we develop a multimodal functional ultrasound and photoacoustic (fUSPA) imaging system - which provides multi-parametric hemodynamics (CBV, CBF, SO2) as well as exogenous molecular imaging of the brain with high spatiotemporal resolutions using a compact head-mountable device. In the following sections, we first present various characterization results of the fUSPA imaging system and its imaging capabilities using phantom, ex vivo and in vivo rat brain studies. Then we show the feasibility of optical and ultrasound contrast agent enhanced in vivo brain imaging. Next, we apply the fUSPA system to study hypercapnia elicited global and regional differences (e.g., arteries vs. veins) in the rat brain using dissected high-resolution and high-sensitivity hemodynamic information. Furthermore, fUSPA imaging for resting-state rats revealed distinct venous dynamics compared to the cortical dynamics. In the future, the high resolution deep brain multiparametric hemodynamic imaging may help gain insights into brain functions in response to various stimuli, mental illnesses [28, 29], brain cancer [30, 31], and transient and complex neurological disorders such as epilepsy [32, 33].

## 2 Results

### 2.1 Functional ultrasound and photoacoustic (fUSPA) imaging system

We developed a multimodal fUSPA imaging system that can perform interleaved UFUS and MSPA imaging of the brain using a compact imaging head. The device tightly integrated a 15 MHz high-frequency ultrasonic transducer (UST) and fiber optic light delivery through lenses such that light and ultrasound fields co-align 7 mm away from the transducer surface (Fig. 1a, Fig. S1, Methods 4.1, and Supplementary Note 1). The optical fiber bundle was coupled to a wavelength-tunable nanosecond pulsed optical parametric oscillator (OPO) laser and the UST was connected to a research ultrasound data acquisition system (DAQ) - Vantage 256 for performing acoustic transmit/receive beamforming and synchronizing triggers with the laser. The Vantage was sequence programmed to continuously obtain fUSPA frames (each consisted of one MSPA and one UFUS block) at 3.3 seconds per frame (Fig. 1b, Fig. S2, and Methods 4.2) to provide the following multiparametric functional cerebral hemodynamic maps: cerebral SO2 map derived from the spectral unmixing of three-wavelength MSPA data (Fig. 1c, Methods 4.3) and CBV and CBF maps obtained from UFUS based multi-plane wave angle compounded data (Fig. 1d, Fig. S2, and Methods 4.4 and 4.5). Additionally, the fUSPA platform can be expanded to facilitate exogenous contrast enhanced brain imaging using a variety of optical and ultrasound contrast agents and respective molecular probes, including MSPA-based exogenous molecular imaging (Fig. 1e) and ULM-based super resolution vasculature imaging (Fig. 1f, Methods 4.6) by intravenous injection of respective optical (e.g., ICG) and ultrasound (e.g., MBs) contrast agents. The integrated imaging of anatomical, functional, and molecular information of the brain at scalable depths and spatial resolutions is a distinct advantage of our fUSPA system, setting it apart from other modalities available.

**Figure 1:**
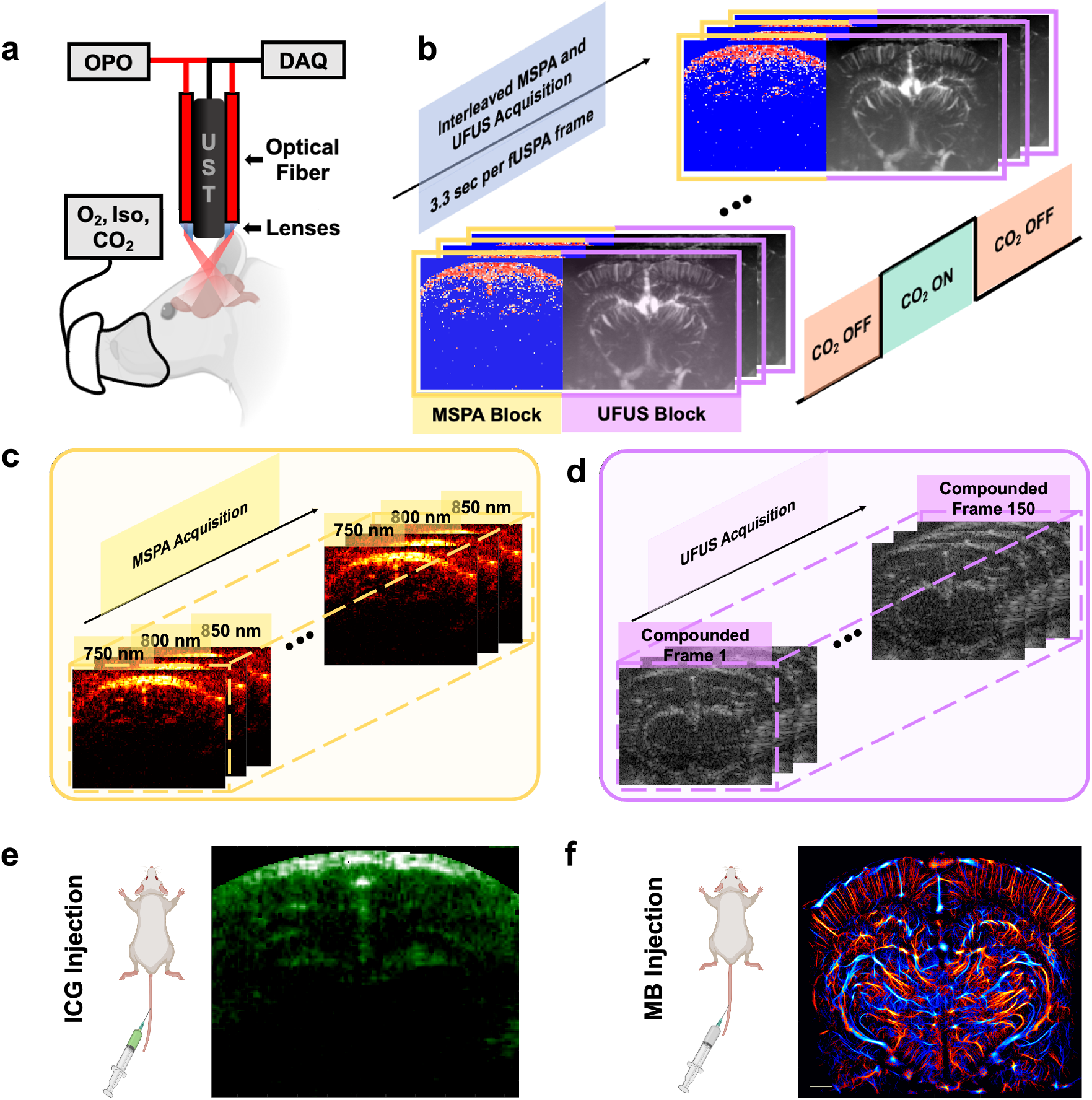
The schematic of the multimodal functional ultrasound and photoacoustic (fUSPA) imaging system and the interleaved imaging sequences. **(a)** The overall schematic of the multimodal fUSPA imaging system using a compact head-mountable imaging device that integrates ultrasound transducer (UST) and optical fiber bundles. The UST probe is connected to the Verasonics data acquisition system (DAQ) to transmit and receive acoustic signals. The fiber optic bundle is connected to the tunable optical parametric oscillator (OPO) laser and delivers light pulses into the brain. The imaging device is water coupled to the transparent chronic cranial window on the rat head. **(b)** The interleaved multimodal fUSPA data acquisition. Each fUSPA frame consists of one multispectral photoacoustic (MSPA) data block and one ultrafast ultrasound (UFUS) data block. The frame rate of fUSPA is around 3.3 frames per second. **(c)** Each MSPA data block consisted of three photoacoustic images acquired at three wavelengths - 750 nm, 800 nm, and 850 nm - in a sequence at 100 ms per PA frame. **(d)** In each UFUS data block, 150 plane-wave angle compounded images were acquired to generate one power Doppler based cerebral blood volume (CBV) image and autocorrelation velocimetry based cerebral blood flow (CBF) map. **(e)** MSPA imaging of intravenously injected photoacoustic contrast agent indocyanine green (ICG) generates vascular perfusion maps. **(f)** UFUS imaging of intravenously injected ultrasound contrast agents, microbubbles (MBs), provides a super-resolved microvasculature imaging using ultrasound localization microscopy principles.

### 2.2 Multiparametric hemodynamic imaging of the brain

Prior to in vivo imaging, multiple characterization steps were carried out to study the performance of the fUSPA imaging system. This includes pulse-echo and PA A-line characterization using a flat metal target, resolution analysis using a wire target phantom, and functional and molecular imaging validation using phantoms consisting of blood flow, SO2, and ICG content (Supplementary Note 2 and Figs. S3 - S5).

We then performed in vivo interleaved UFUS and MSPA imaging of anesthetized rat brain by coupling the compact fUSPA imaging head to the chronic cranial window that is transparent to both light and ultrasound (see Methods 4.2 and 4.10). Fig. 2 shows multiparametric CBV, CBF, and SO2 maps from three distinct coronal locations (Bregma -5.5, -4.0, and -2.7) of the rat brain overlaid with corresponding atlases [34]. UFUS data was processed to display both grayscale power Doppler intensity maps representing CBV changes (Fig. 2b, Methods 4.4) and the color coded ultrasound velocimetry maps quantifying CBF (Fig. 2c, Methods 4.5), with red and blue colors denoting descending and ascending blood flow directions in the microvasculature. The SO2 maps for the three coronal brain locations (Fig. 2d, Methods 4.3) are obtained from the linear spectral unmixing of MSPA frames at wavelengths of 750 nm, 800 nm, and 850 nm (Fig. S6) overlaid with the corresponding CBV maps. Unlike ultrasound based CBV and CBF maps, the SO2 information was limited to approximately 6 mm depth, primarily due to limited light illumination through the cranial window and optical attenuation within the brain tissue. This limitation is consistent with other in vivo B-mode PA brain imaging studies [25, 26] as well as our ex vivo studies (Fig. S7). Furthermore, the significantly higher SO2 values obtained in vivo (Fig. 2d) compared to ex vivo imaging (Fig. S7) validated the capability of MSPA for monitoring cerebral SO2 changes. Additionally, as shown in Figs. 2e-2g, it is possible to generate volumetric (3D) maps of CBF, CBV and SO2 by combining respective 2D (B-mode) cross-sectional images obtained from translating the compact fUSPA imaging head along the anterior-posterior axis (300 *µ*m step size). These results demonstrated that the fUSPA imaging system can provide brain-wide multiparametric hemodynamic information in vivo with high spatial resolution and sensitivity.

**Figure 2:**
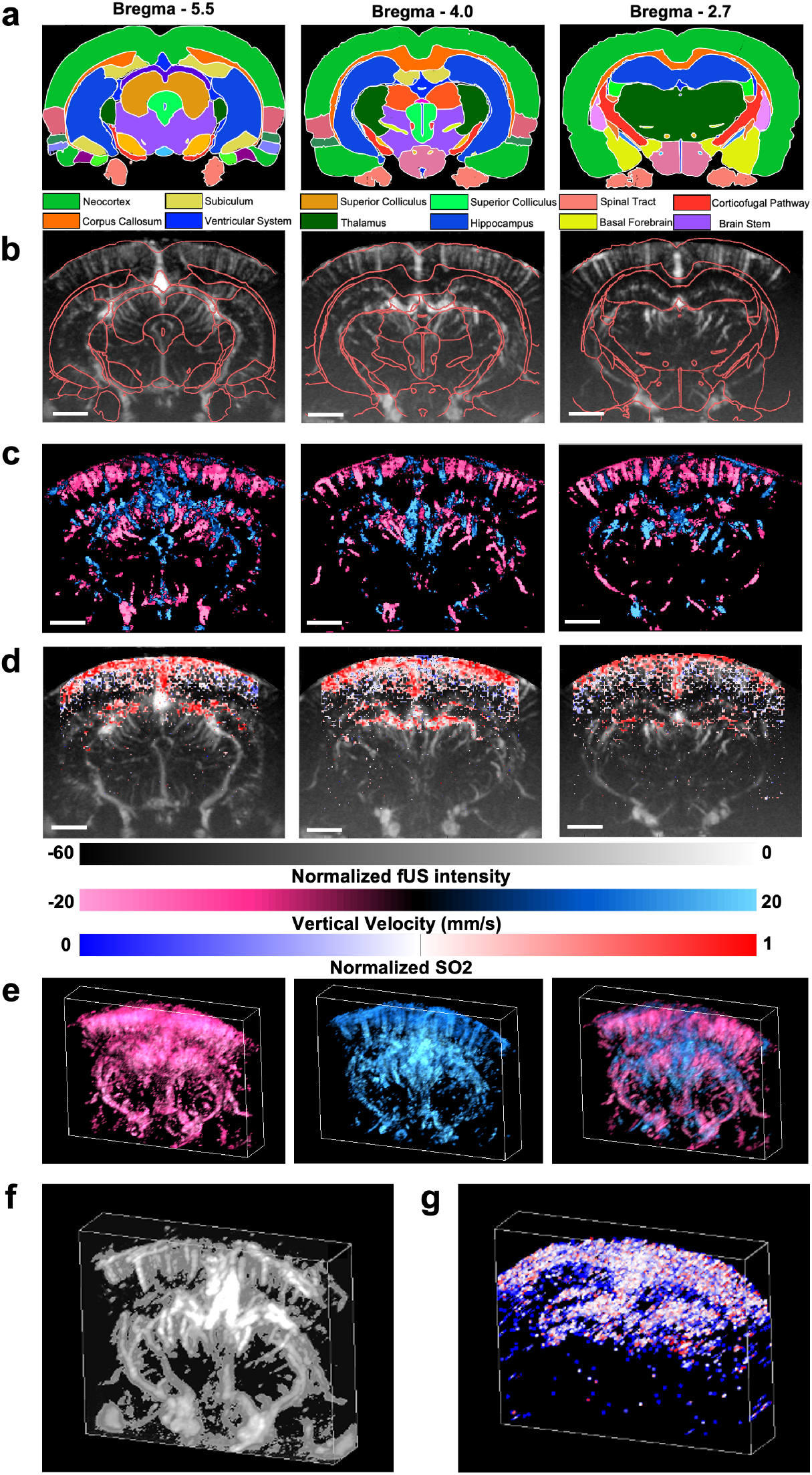
In vivo ultrafast ultrasound (UFUS) and multispectral photoacoustic (MSPA) imaging of rat brain provided multiparametric hemodynamic brain activity maps. **(a)** The coronal rat brain atlas maps at Bregma - 5.5, Bregma -4.0 and Bregma - 2.7 locations [34] with different brain regions in different colors. **(b)** Three CBV images and **(c)** three CBF images were generated from the UFUS data acquired from the three Bregma locations, and overlaid with the respective atlas contours in red color. In the CBF map, the vertical direction of the measured flow is color coded, where red color denotes descending flow (downward) and blue color denotes ascending flow (upward). **(d)** The MSPA data based unmixed oxygen saturation (SO2) maps at the three Bregma locations are co-registered onto the grayscale CBV images. Volumetric maps of **(e)** CBF **(f)** CBV and **(g)** SO2 were generated from the B-mode UFUS and MSPA data acquired with linear scanning of the imaging device. In (e) red and blue color denotes descending and ascending flow maps and the combined CBF map on the right. CBF: cerebral blood flow; CBV: cerebral blood volume. Scale bars represent 2 mm.

### 2.3 Exogenous contrast enhanced brain imaging

We studied the exogenous contrast enhanced brain imaging capabilities of the fUSPA system by intravenous injections of US and PA contrast agents in conjugation with UFUS and MSPA imaging, respectively. By localizing and tracking the intravenously injected MBs across multiple compounded frames acquired during the 10-minute UFUS imaging period, we generated a super resolved (10 *µ*m *×* 10 *µ*m) ULM images of brain vasculature covering the whole cross section of the rat brain (Fig. 3a) [35, 36]. The tracking of microbubbles also allowed us to determine the vertical direction of flow (Fig. 3b) and calculate the mean flow speed (Fig. 3c) (Methods 4.6).

**Figure 3:**
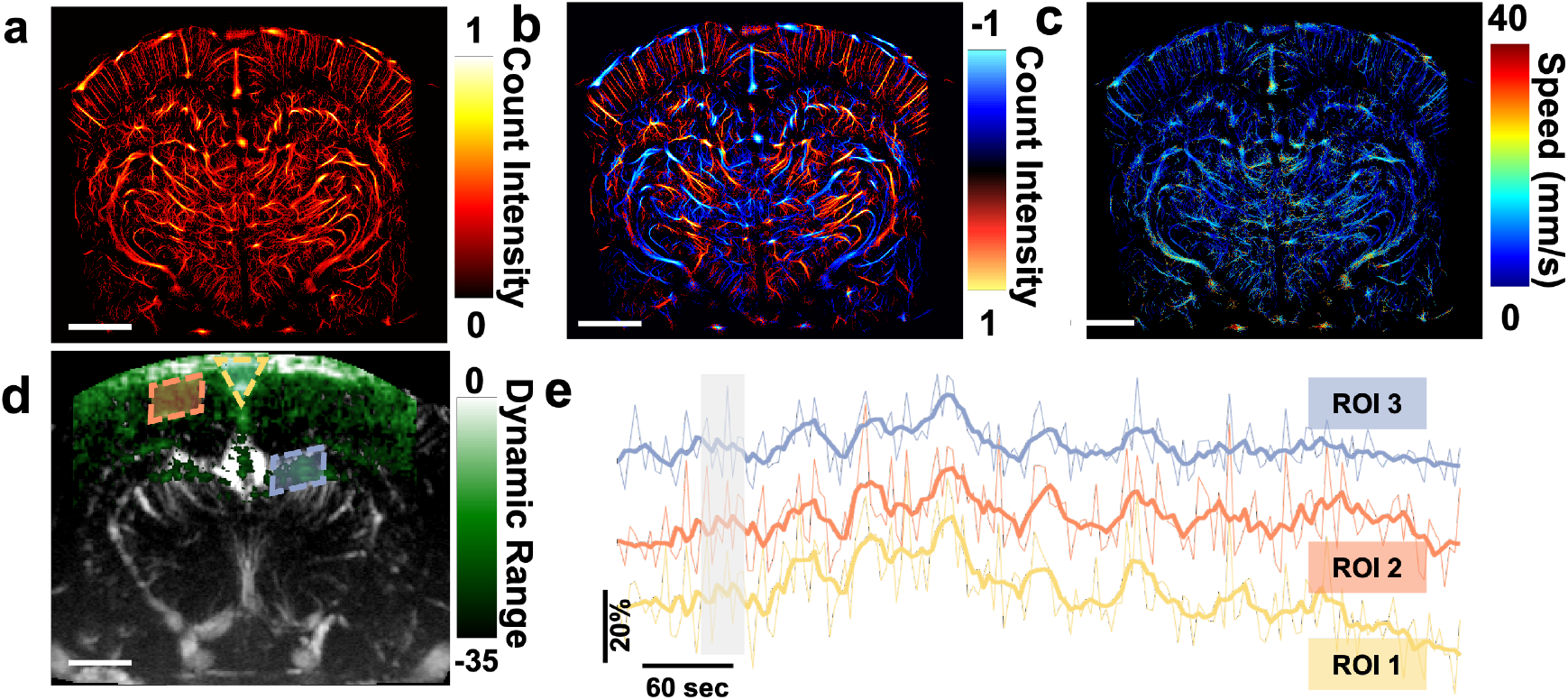
Exogenous contrast agent enhanced fUSPA imaging of rat brain. **(a-c)** Super-resolution ultrasound localization microscopy (ULM) imaging of rat brain microvasculature using intravenous injection of microbubbles (MBs). ULM images based on **(a)** MB count intensity and **(a-c)** Super-resolution ultrasound localization microscopy (ULM) imaging of rat brain microvasculature using intravenous injection of microbubbles (MBs). ULM images based on **(a)** MB count intensity and **(b)** MB count intensity color-coded with vertical flow direction; red and blue color denote the bubbles moving downward and upward respectively. **(c)** By calculating the displacement and corresponding frame rate, an averaged blood speed map is generated. **(d)** MSPA based vascular perfusion imaging using intravenously injected small molecule photoacoustic contrast agent, indocyanine green (ICG) overlaid with UFUS based CBV map at around Bregma -4.5. **(e)** The plots of ICG time activity averaged over selected ROIs showed the relative ICG changes during pre-injection, injection, and post-injection periods. The gray shaded area represents the time during ICG injection. The bolded lines represent the smoothed average ICG changes in the ROI. MSPA: multispectral photoacoustic; CBV: cerebral blood volume; ROI: region-of-interest. Scale bar of the images represents 2 mm.

Next, we studied exogenous photoacoustic molecular imaging capabilities of the fUSPA system by intravenously injecting ICG molecules, while continuously acquiring MSPA data at 750 nm, 800 nm, and 850 nm wavelengths for a total time period of 10 minutes (see Methods 4.11). Linear spectral unmixing of each set of the three-wavelength MSPA frames provided a map of ICG molecular distribution inside the brain (Fig. 3d). The comparison of the three-wavelength MSPA frame sets acquired at pre- and post-ICG injection shows that the overall brain PA intensity post-injection increased by 2 dB, 5 dB, and 3 dB for 750 nm, 800 nm, and 850 nm wavelengths respectively (Fig. S8). The strongest difference observed at 800 nm corresponded well with the peak optical absorbance of ICG in plasma at 800 nm wavelength [37]. Additionally, unmixing of all the acquired temporal MSPA frames provided time-stamped unmixed ICG images, which were used to track the temporal dynamics of ICG molecule accumulation and its subsequent clearance during the post-injection phase (Fig. 3e, Methods 4.12). All three region-of-interests (ROIs) inside the brain - sagittal sinus (ROI 1), cortical vessels (ROI 2), and hippocampal vessels at *∼* 3.5 mm depth (ROI 3) - showed peak uptake of ICG around 2 minutes post-injection, which aligned well with the 1.5 - 3 minute ICG half-life in rats [37].

### 2.4 Studying stimulation-evoked multiparametric hemodynamic changes at single-vessel resolution

High-resolution, high-sensitivity, and multiparametric imaging capabilities of the fUSPA system were then leveraged to study stimulation evoked functional hemodynamic changes of the brain at single vessel resolution. Hypercapnia, a widely-adopted carbon dioxide (CO_2_) stimulation strategy to assess the vascular reactivity in various diseases [38–40], was chosen as it elicits global vasodilation as well as elevating oxygen saturation inside brain and other organs [41–43].

With the animal under anesthesia (1 - 1.5% isoflurane mixture with O_2_), we first conducted continuous interleaved UFUS and MSPA imaging at Bregma -5.8 (Fig. 4a) during the 8-minute CO_2_ stimulation paradigm (2 minutes of baseline, 3 minutes of 5% CO_2_ stimulation, and 3 minutes of post-stimulation, see Methods 4.11). Next, using intravenous MB injection, ULM imaging of the same brain location was obtained to map microvascular anatomy and vertical flow directions (Fig. 4b). While UFUS data was processed to provide both CBV and CBF maps as shown in Fig. 4c and 4e respectively, the spectral unmixing of MSPA data provided SO2 maps (Fig. 4d) of the brain during the stimulation. In Fig. 4b ULM and Fig. 4e CBF images, the red and blue colors denote downward and upward flows, and in the cortical regions these colors represent arterioles and venules, respectively.

**Figure 4:**
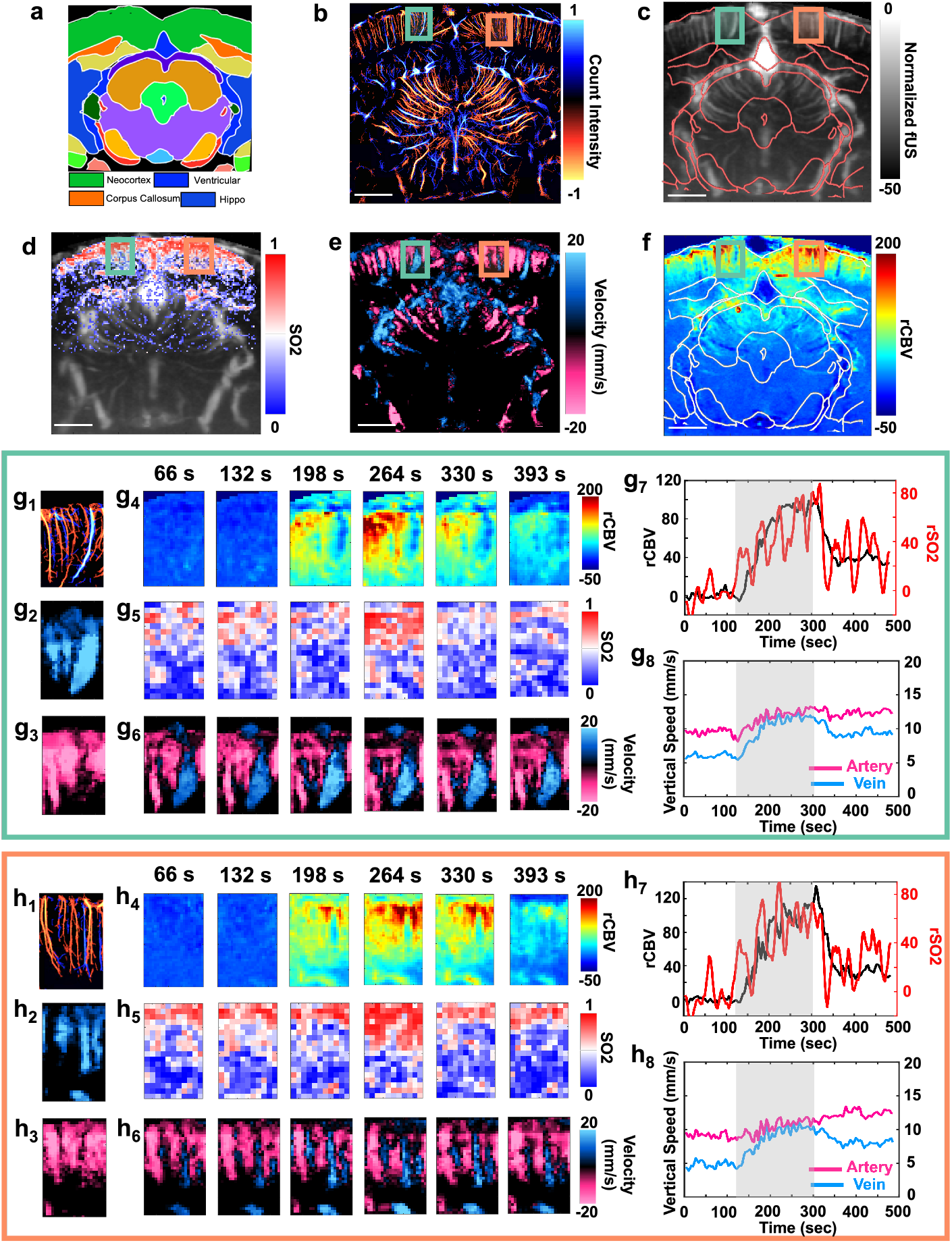
Multiparametric fUSPA imaging revealed regional differences in cerebral vascular reactivity during hypercapnia (CO2) stimulation. **(a)** A coronal section map at Bregma -5.8 location of the rat brain atlas [34] and **(b)** ULM based super resolved microvasculature imaging at the Bregma -5.8 location. Red and blue colors denote downward and upward flows. The color bar represents the normalized count intensity. **(c)** The UFUS based CBV image overlaid with the atlas from (a), and **(d)** the SO2 map obtained from MSPA imaging overlaid onto the corresponding CBV map. **(e)** The CBF map was generated from the velocity autocorrelation of UFUS frames, where red and blue denotes downward and upward blood flows. In the cortex region, the descending vessels were arterioles and the ascending vessels were venules. **(f)** The relative CBV (rCBV) change map during hypercapnia (t = 260 second). Scale bars represent 2 mm. Two ROIs in the cortex of images in (b) to (f) were enlarged and shown in **(g)** and **(h)** to study regional differences in the hemodynamic responses. In (g) and (h), the first subfigures (g1, h1) display ULM based microvascular map inside the corresponding ROI, with a scale bar of 0.5 mm. The second (g2, h2) and third (g3, h3) subfigures display the vessels with ascending and descending flows obtained from UFUS based CBF maps of the two ROIs, respectively. Subfigures display (g4, h4) the rCBV, (g5, h5) the SO2, and (g6, h6) the CBF dynamics inside the two ROIs for selected time points during the hypercapnia stimulation. The (g7, h7) subfigures plot the averaged rCBV (black line) as well as the averaged rSO2 (red line) changes in the respective ROIs during the hypercapnia stimulation. The (g8, h8) subfigures display differential vertical CBF between arteries and veins, determined from the flow direction, during the hypercapnia stimulation. The gray shaded area in g7, g8, h7, h8 represent the time period with CO2 on. CBV: cerebral blood volume; rCBV: relative cerebral blood volume; CBF: cerebral blood flow; ULM: ultrasound localization microscopy; ROI: region-of-interest; SO2: oxygen saturation; rSO2: relative oxygen saturation; MSPA: multispectral photoacoustic.

We further calculated the relative changes in CBV and SO2 maps – referred as rCBV and rSO2 respectively - as well as CBF maps during the hypercapnia stimulation paradigm (Methods 4.12). Although the stimulation increased the CBV, CBF, and cerebral SO2 globally (Supplementary Movie S1, Movie S2 and Movie S3), differential temporal changes were observed between cortical vessels, as shown in the representative rCBV map. To further study the regional differences in the stimulation evoked hemodynamic changes, we analyzed CBV, CBF, and SO2 information in the ROI 1 and ROI 2 (green and orange boxes) as shown in Figs. 4g-h. During the hypercapnia stimulation, the rCBV showed *∼*100% increase in ROI 1 (Fig. 4g_4_ and Fig. 4g_7_) and *∼*120% increase in ROI 2 (Fig. 4h_4_ and Fig. 4h_7_), and overall rSO2 synergistically increased *∼*80% for ROI 1 (Fig. 4g_5_ and Fig. 4g_7_) and *∼*60% for ROI 2 (Fig. 4h_5_ and Fig. 4h_7_). Based on the high-resolution ULM microvascular map (Fig. 4g_1_), we delineated ROIs around the singular vein and artery in ROI 1 (Fig. S9a and Fig. S9b) and observed that rCBV change in the vein (*∼*15% increase) was much less compared to the artery (*∼*160% increase) at the peak stimulation time point (around t = 260 seconds). In contrast to the CBV changes, the analysis of CBF dynamics (Fig. 4g_2_-4g_3_, 4g_6_, 4h_2_-4h_3_, 4h_6_, S9d-f) showed a different case, where the arterial CBF demonstrated lower relative increase (*∼*31.6% increase for ROI 1, from *∼*9.5 mm/s to *∼*12.5 mm/s - Fig. 4g_8_, and *∼*25.6% increase for ROI 2, from *∼*9 mm/s to *∼*11.3 mm/s - Fig. 4h_8_) comparing to that of the veins (*∼*91.7% increase for ROI 1, from *∼*6 mm/s to *∼*11.5 mm/s - Fig. 4g_8_, and *∼*122.2% increase for ROI 2, from *∼*4.5 mm/s to *∼*10.0 mm/s - Fig. 4h_8_). These differential CBF changes also held valid when drawing ROIs around a singular arteriole and venule (Fig. S9d-f).

Surprisingly, we observed strong reductions of rCBV during hypercapnia in the superior sagittal sinus (SSS, Fig. S10) and the ventricle (Fig. S11) regions of the brain. Although both SSS and ventricle regions exhibited strong CBV intensity (Fig. S10a, Fig. S11a), the rCBV calculated during hypercapnia showed a decrease of *∼*-40% for SSS (Fig. S10e and S10g) and *∼*-17% for ventricle (Fig. S11e and S11g) regions. On the other hand, the rSO2 for SSS showed an increase of around 50% (Fig. S10f and S10g), but minimal to no change was shown for the ventricle region (Fig. S11f and S11g). This is likely because SSS consists of large veins that drain significant amount of blood, whereas the ventricle region primarily consists of the complex capillary bed (see enlarged ULM image of the ventricle in Fig. S11d) in the choroid plexus that produces cerebrospinal fluid (CSF), which does not absorb sufficient light to generate photoacoustic contrast.

### 2.5 Correlation between cortex and vein temporal dynamics during the resting and stimulation states

The fUSPA system was further employed to study the hemodynamic changes in the sagittal section near the midline of the rat brain. The CBV map and the corresponding rCBV map during hypercapnia stimulation are shown in Fig. 5a and Fig. 5b, respectively, and the ULM based vertical flow map is shown in Fig. 5c. Similar to the coronal plane imaging results, the overall cortex (green shaded area - ROI 1 in Fig. 5a) showed an increase in rCBV, while the veins (example vein shown as purple shaded ROI 2 area within the white solid ROI 3 box of Fig. 5a-c) exhibited minimal rCBV change (Fig. 5b). The veins in the ROI 2 were identified using ULM vertical flow map (Fig. 5c), and corresponding lower SO2 value shown in Fig. S12. Similar to the previous observation in Fig. 4, although the ventricle region (white dashed inlet - ROI 4 in Fig. 5a-c) showed strong CBV intensity (Fig. 5a), negative rCBV and little SO2 content (Fig. S12) were found during hypercapnia stimulation (Fig. 5b).

**Figure 5:**
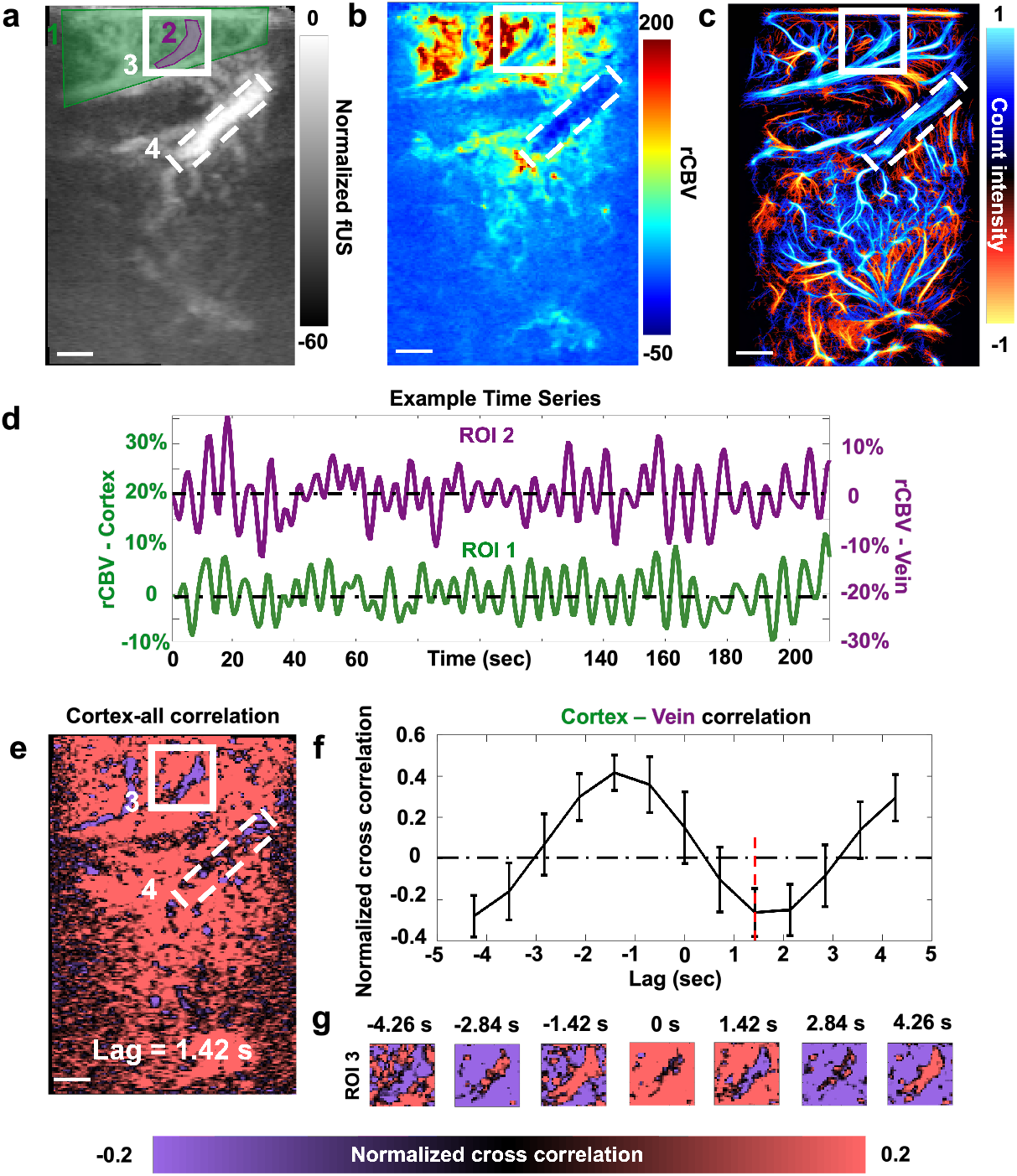
Correlation between the venous and cortical hemodynamics revealed by fUSPA. Data in this figure are acquired with the fSUPA device positioned to image sagittal section of the brain at around the midline. **(a)** The CBV map. The cortical CBV dynamics were averaged across the green shaded area (ROI 1) and the venous CBV dynamics were extracted from the purple shaded area (ROI 2) within the white solid ROI (ROI 3). The ventricle region is delineated with white dashed ROI (ROI 4). **(b)** The rCBV change map during the hypercapnia stimulation. **(c)** The corresponding ULM image showed the vertical flow direction in the sagittal plane. **(d)** The time activity of the averaged cortical rCBV (in green, from ROI 1) and averaged venous rCBV (in purple, from ROI 2). **(e)** The cross-correlation map between the temporal dynamics of averaged cortical CBV and the temporal dynamics of each voxel in the CBV image, with the strongest coupling at lag = 1.42 sec is displayed in (e). **(f)** The cross-correlation plot between the cortex (ROI 1) and the venous (ROI 2) CBV signals. The strongest decorrelation was highlighted with the red dashed line at lag =1.42 seconds. **(g)** The enlarged cross-correlation map of ROI 3 at different lags. The scale bars represent 1 mm. ROI: region-of-interest; CBV: cerebral blood volume.

fUSPA also allowed us to study the correlation between the veins (ROI 2) and overall cortical hemodynamics (ROI 1) during the resting state by performing continuous UFUS data acquisition for 5 minutes on the anesthetized (1 - 1.5% isoflurane) animal without stimulation. In the rCBV time series maps derived from continuous UFUS data, the averaged rCBV intensity for cortical (ROI 1) and venous (ROI 2) regions plotted as a function of time are plotted in Fig. 5d, and demonstrated a clear out-of-phase dynamics (Fig. 5d). To further analyze this temporal pattern across the imaging plane, we calculated the cross-correlation between the averaged cortical (ROI 1) dynamics and all voxels in the CBV map, denoted as cortex-all correlation (full lag maps in Fig. S13, example Lag = 1.42 sec in Fig. 5e, Methods 4.14) and the cross-correlation between the cortical and venous dynamics, denoted as cortex-vein correlation (Fig. 5f, Methods 4.14). Interestingly, although the majority of the voxels showed positive correlations with the cortical regions (Fig. 5e, S13), the venous dynamics, however, were clearly anti-correlated with other regions at all Lags (Fig. 5e, 5g, S13), with cortex-vein correlation showed a positive peak at Lag = -1.42 sec (R = 0.37, Fig. 5f) and a negative peak at Lag = +1.42 sec (R = -0.26, red dashed line in Fig. 5f). This lag observed in the temporal pattern aligns with previous measurements of transit time in rodents [44]. However, the slow wave cortex-vein coupling, as obtained using high spatiotemporal resolution fUSPA, distinguishes itself from other imaging modalities.

## 3 Discussions

In this article, we present a novel in vivo multimodal fUSPA imaging system to quantitatively map multiparametric hemodynamic and molecular information of deep brain regions of rats with high spatiotemporal resolutions. Effective integration of hardware and software components of fUSPA enabled interleaved UFUS and MSPA data acquisitions in real-time using a single head-mountable imaging device. UFUS data processing provided both power Doppler based CBV mapping and ultrasound velocimetry based CBF mapping at single vessel resolution. The spectral unmixing of MSPA data provided cerebral SO2 maps. Furthermore, through the intravenous administration of FDA approved ultrasound and optical contrast agents, MBs and ICG respectively, fUSPA system allowed ULM based super-resolved microvasculature imaging and ICG molecule enhanced vascular perfusion imaging of the brain. In total, the fUSPA imaging system can provide the following six brain maps: CBV, CBF, SO2, ICG vascular perfusion, super-resolved microvascular anatomy, and axial blood velocity. Such comprehensive mapping of individual and dissected hemodynamic parameters provides fUSPA imaging distinct advantages over the BOLD fMRI, which represents a mixed effect from CBV, CBF, and cerebral metabolic rate of oxygen consumption (CMRO_2_) that confound interpretations [9, 10, 45].

Using high-resolution multiparametric hemodynamic imaging capabilities of the fUSPA system, we investigated brain wide changes in hemodynamics elicited by hypercapnia stimulation. Traditionally, fMRI [46], optical imaging [47], or a combination of these modalities [48] are used to study the hypercapnia effect on different brain regions or global cerebrovascular reactivity. However, using fUSPA, which offers superior imaging depth compared to pure optical imaging and improved spatiotemporal resolution and high sensitivity compared to fMRI, we could identify individual vessel-specific effects of hypercapnia that other modalities often miss. As hypercapnia is a known vasodilator and increases tissue oxygenation [43], our results confirmed a global and synergistic increase in CBV, CBF, and SO2 (Supplementary Movies S1-3), but revealed regional differences - e.g., the SSS region exhibited a decrease in rCBV but increase in rSO2 values during the hypercapnia stimulation. Further leveraging the high-resolution of the fUSPA system, single vessel analyses were conducted and differential hemodynamic response during stimulation was found between the cortical arteries and veins. Strikingly, cortical veins exhibited minimal rCBV increase but significant rCBF increase compared to the arteries. The revealed rCBV differences between arteries and veins were similar to other sensory stimulation [49–51], and the observed differential CBF and CBV changes agreed with previous hypercapnia studies based on Doppler ultrasound [52] and positron emission tomography [53], respectively. However, our results show the first evidence of multiparametric hemodynamic changes at a single vessel spatial resolution, making it suitable to study microvascular cerebral metabolisms [54] and brain injuries [55].

The advantage of deep brain single vessel multiparametric information further extended to imaging the resting state brain in the sagittal plane. The analysis of continuous UFUS data during the rat resting state showed the venous dynamics were out-of-phase from the overall cortical dynamics with approximately 1.42 seconds delay. Since CBV is directly proportional to the volume changes of the moving scatterers (where hemoglobins are the main scatterers in the vessels), this consistent lag between venous (ROI 2 in Fig. 5) and cortical dynamics (ROI 1 in Fig. 5) suggested a reproducible temporal pattern between the venous blood volume and overall CBV, which further agreed with previous findings using optical imaging [44] and fMRI [56]. However, optical imaging is not capable of mapping deep brain venous dynamics, while fMRI could not directly assess the venous rCBV change [56], setting fUSPA as a unique device for studying the temporal patterns of the single vessel in the deep brain, which is often assessed in sleep [57], and resting state studies [58]. Furthermore, fUSPA provided the multiparametric ULM based vertical speed map and MSPA based SO2 map that helped us confidently segment out the venous area, which was not achievable with fMRI alone. Remarkably, both ventricle and venous regions showed decreased rCBV during hypercapnia stimulation (Fig. 4, Fig. S11b, and Fig. 5b) and decorrelation with the cortical dynamics during the resting state (Fig. S13). While the CBV intensity was strong in the whole ventricular region (Fig. S11a and Fig. 5a), the correlation maps (Fig. 5e-g) and the SO2 map (Fig. S11 and Fig. S12) showed only low and scattered values. Considering the web-like microvascular structures in the ventricle mapped by ULM, we hypothesize that the strong CBV signal was primarily from the fast flowing blood in the choroid plexus capillaries that resided in the ventricle [59, 60]. Many prior studies reported that the venous dynamics were coupled and correlated with the CSF signal [61–63], especially during deep/forced breathing. Therefore, given that the choroid plexus is responsible for the production of the majority of CSF [64], we reasonably hypothesize that the venous-ventricle correlation measured by CBV mapping was related during isoflurane anesthesia. However, this hypothesis warrants further studies.

The fUSPA imaging system reported here does have some limitations. Firstly, although our light delivery is compactly integrated into the transducer device, it still requires a few millimeters of coupling medium (working distance) for proper illumination. This distance not only hinders the system’s ability for mobile animal imaging but also increases the data transfer size, leading to a reduction in temporal resolution. To address this, future improvements can employ a transparent ultrasound transducer (TUT) array [65], which would allow seamless integration of multimodal UFUS and MSPA with minimal coupling distance. Further, a multi-angle plane wave light illumination through the TUT array can help improve the depth of MSPA imaging. Additionally, the vascular compartment segmentation, specifically distinguishing between veins and arteries, is applicable only for cortical vessels since the ULM and CBF map would only be able to encode the vertical flow direction (upwards/ascending or downwards/descending) relative to the cortical surface. This limitation may be mitigated by generating 3D ULM images of the entire brain using a 2D matrix probe [66], and then encode three dimensional flow direction to further understand the blood flowing pathway [67].

Despite the above mentioned limitations, our presented novel multimodal fUSPA system is the first of its kind that enables deep brain comprehensive hemodynamic mapping (CBV, CBF, and cerebral SO2) and exogenous contrast enhanced brain imaging with high resolution and sensitivity. The unique multiparametric capabilities of fUSPA will help advance our understanding of complex brain functions, thereby paving the way for better insights into various neurodiseases, such as brain cancer, sleep disorders, and Alzheimer’s disease.

## 4 Methods

### 4.1 fUSPA imaging device

To achieve a compact fUSPA imaging head, we designed a light-weight portable holder that accommodated ultrasound transducer (UST) in the center and fiber optic light delivery through acrylic lenses from either side of the UST to focus light into the ultrasound detection zone (Figs. S1a and S1b). The high-frequency UST (Vermon, Tours, France, 15 MHz, 40% fractional bandwidth, 128 elements with 0.1 mm pitch, 1.5 mm elevation, and 6 mm elevational focus) was custom designed, and the overall size of UST was 5 mm *×* 20 mm *×* 25.4 mm (width *×* length *×* height) with a weight of around 5 grams. The high-frequency UST was used for both UFUS plane wave imaging and PA detection. The fiber optic delivery consisted of four rectangular fiber bundles (Fiberoptic Systems Inc, Simi Valley, CA, USA), each with a size of 2 mm *×* 5 mm, cased inside a 2.5 mm *×* 6 mm stainless steel tubing, and inserted into the holder on each side (Figs. S1a-b). Two 50-degree lenses machined from polished acrylic sheets were attached to the front of the optical fiber bundles to refract the optical beams such that they focus at around 6-7 mm away from the transducer surface, and allow co-alignment with the ultrasound field inside the tissue medium (Figs. S1c-d and see Supplementary Note 1 for the calculations). The focused beam at *∼* 7 mm was measured to cover an area of around 3 *×* 26 mm^2^, which was similar to the theoretical calculation of 2.1 *×* 20 mm^2^. For animal imaging, the laser energy was controlled by tuning the Q-switch delay, and the output laser energy after the lenses were measured by an energy meter (PE50BF-DFH-C, Ophir-Spiricon, LLC, North Logan, UT, USA) to be less than 15 mJ/pulse, which gave an optical fluence below the American National Standards Institute (ANSI) safety maximum permissible exposure (MPE) limit of 20 mJ/cm^2^ [68].

### 4.2 Interleaved UFUS and MSPA data acquisition

A Vantage 256 ultrasonic research system (Verasonics Inc., Kirkland, WA, USA) was custom programmed to perform interleaved MSPA and UFUS imaging, including transmitting ultrafast plane wave angle compounded US imaging, receiving US echoes and PA pressure waves, and online/offline reconstruction using pixel-based beamforming. The schematic of the combined fUSPA imaging sequence for interleaved MSPA and UFUS data acquisition is shown in Fig. S2 and described in detail below.

Each fUSPA frame data was acquired in 3.3 seconds and consisted of 1.9 seconds for MSPA data, *∼* 1.2 seconds for UFUS data acquisition, and *∼* 0.2 seconds for trigger synchronization (Fig. S2a). As shown in Fig. S2b, each MSPA data block was composed of 16 dual-modality B-mode US and PA (USPA) frames, with each B-mode US and B-mode PA frame acquired every 100 ms (Fig. S2b), limited by the 10 Hz pulse repetition rate of the tunable optical parametric oscillator (OPO) laser (Phocus Mobile, Opotek Inc., Carlsbad, CA, USA; 5-7 ns pulse length, peak pulse energy of 100 mJ at 750 nm, with tuning range of 690 nm to 950 nm and 1200 nm to 2600 nm). While B-mode US frame shows mechanical impedance contrast based anatomical information of the tissue, B-mode PA frame displays optical absorption contrast based molecular information of the tissue. Since both B-mode US and PA frames are acquired using the same UST probe, a co-registered US+PA (USPA) frame with overlaid anatomical and optical contrasts is helpful to understand the origin of PA signals [17, 65]. After acquiring each USPA frame data, the Vantage was set to idle to wait for the laser trigger pulse (Fig. S2b), which was achieved by using an external function generator (SDG6022X, Siglent Technologies NA, Solon, OH, USA) as the master (similar to the Verasonics recommended strategy [69]) to minimize jitters for PA signal acquisition at other wavelengths. The fast-tuning feature of the OPO laser allowed us to tune the wavelength for every 100 ms, enabling real-time MSPA imaging. Three wavelengths - 750 nm, 800 nm, and 850 nm - were chosen for MSPA imaging, and 15 MSPA frames (5 sets of 750 nm, 800 nm, and 850 nm PA frames) were acquired to achieve sufficient signal-to-noise ratio. One additional MSPA frame was acquired due to potential frame loss during laser trigger synchronization, giving a total of 16 MSPA frames. With the 10 Hz laser pulse repetition frequency, 16 MSPA frames (along with respective US frames) took 1.6 seconds, and 0.3 seconds were taken to transfer and save the data, which totaled up to be *∼* 1.9 seconds shown in Fig. S2b.

Then UFUS data was acquired right after the MSPA data transfer. The timing diagram of the UFUS data acquisition sequence block is displayed in Fig. S2c. During one UFUS acquisition, 4 half cycles of 11 volts were used to excite the ultrasound transducer [70] with seven (7) planar ultrasonic waves tilted at angles varying from -6 to 6 degrees (-6*^◦^*, -4*^◦^*, -2*^◦^*, 0*^◦^*, 2*^◦^*, 4*^◦^*, 6*^◦^*, with 75 *µ*s between each plane wave angle, which are coherently summed to generate one compounded US frame every 525 *µ*s. 150 of such compounded frames (US*_C_*_1_ - US*_C_*_150_ as shown in Fig. S2c) were used to generate one power Doppler based CBV map in 78.75 ms [11] and an additional 1.1 seconds was required for data transfer (Fig. S2c). Then the system was set to idle and awaited the trigger for synchronization and ready for the next fUSPA frame acquisition.

### 4.3 MSPA based molecular imaging

We used linear unmixing based on non-negative least squares curve fitting to unmix different molecular distributions (HbO, HbD, and ICG) inside the tissue. The input matrix for curve fitting was initially obtained from omlc.org and normalized based on the measured OPO laser pulse energy differences across the three wavelengths (750 nm, 800 nm, and 850 nm). As discussed in the above section and Fig. S2b, each MSPA data block of 1.6 seconds duration consisted of 15 PA images, acquired in 5 sets of 3 PA images (one PA image per each of the three wavelengths). The resulting 5 PA images for each of the three wavelengths are averaged to improve the signal-to-noise (SNR), and averaged three-wavelength PA images are used in the spectral unmixing to generate unmixed images of HbO, HbD and ICG. While the unmixed ICG map was directly displayed after log-compression (e.g., Fig. 3d), the SO2 map was generated from the unmixed HbD and HbO images, by calculating the SO2 = HbO/(HbO + HbD) and normalization across all the SO2 values for each pixel (e.g., Fig. 2d).

### 4.4 UFUS based CBV imaging

For each UFUS data block, we generated one power Doppler map as a CBV proxy using similar processing as the existing literature. We coherently compounded the in phase/quadrature (IQ) data and then applied singular value decomposition (SVD) filter [71] to remove the tissue motion (first 9 ranks) from the 150 compounded frames while preserving the signal components associated with the moving blood signal. As red blood cells are the main scatterers in the blood, power Doppler signal intensity is considered to be directly proportional to the fractional moving CBV [11]. Hence, the power Doppler intensity map is considered as the CBV intensity map.

### 4.5 UFUS based CBF imaging

To study the dynamic CBF changes, a velocimetry map was derived from the acquired UFUS data. The processing was similar to the previously reported method by Tang et al. [23]. In summary, the algorithm fits autocorrelation functions using nonlinear regression to both the positive and negative component of the IQ temporal signal from the 150 compounded frames to solve the z direction velocity.

In detail, each UFUS compounded frame data was spatiotemporally filtered using SVD by rejecting components corresponding to the first 10 singular values, followed by a fourth-order Butterworth 70 Hz high pass filter to generate the moving blood IQ signal. Then the IQ signal was subjected to directional filter to obtain positive and negative signal components based on the frequency spectrum, as a clear separation of the opposing frequency components is observed based on the blood flow direction relative to the UST [72]. The positive and negative signals were only considered valid for later processing if their individual positive and negative frequency powers were greater than 0.2 and 0.25 of the total frequency power, respectively. In addition, to remove the effect of noise, only the signals with the first temporal autocorrelation greater than 0.2 were used in subsequent processing [23]. Then a first-order temporal autocorrelation function was used to separately fit the filtered signals to solve the z direction velocity as velocity maps as well as their corresponding flow direction information. The velocity maps generated by this fitting procedure were found to agree with the vertical speed obtained from the ULM maps [23].

### 4.6 ULM imaging

ULM was acquired using the same fUSPA system with the help of intravenous injection of ultrasound contrast agents - MBs into rats. The ULM data acquisition was similar to the above described UFUS data acquisition, except with minor changes in the time between the angles for plane wave imaging and the number of compounded frames. In particular, ULM imaging was performed using plane waves transmitted at 7 angles (-6*^◦^*, -4*^◦^*, -2*^◦^*, 0*^◦^*, 2*^◦^*, 4*^◦^*, 6*^◦^*) with 200 *µ*s between each angle to generate compounded images with a frame rate of *∼* 700 Hz. 400 compounded images (560 ms duration) were acquired for each sequence and then *∼*2 seconds were used to transfer the data to the host, which gave a sequence duration of *∼*2.6 seconds. 300 such sequences were acquired, thereby giving a total ULM accumulation time of 168 seconds with a total acquisition time of around 9 minutes. The ULM images were obtained from beamforming of the raw RF data using the Verasonics system to generate the IQ data. The IQ dataset was filtered using the spatiotemporal SVD filter [71], where the first few singular values were removed to isolate MB signals from the surrounding tissue signals. ULM MB localization procedure was similar to [35, 73]. In brief, MBs were localized to sub-pixel resolution using spline-based interpolation. Thereafter, we used the Hungarian algorithm to form tracks [74] with a minimum duration of 100 seconds. The tracks were then accumulated to form the vasculature map [73]. The displacement of MBs between two consecutive frames was used to estimate the speed of tracked MBs, and the speed of all MB traces during the ULM session was averaged to form the flow speed map [75].

### 4.7 Resolution wire targets

The resolution of the US and PA was characterized by imaging 50 *µ*m diameter micro metal wire targets embedded at different depths inside a tissue-mimicking phantom (Fig. S4). To generate strong ultrasound and photoacoustic contrasts, three 50 *µ*m diameter micro metal wires (W1 - W3 as shown in Fig. S4a) were dyed with India ink, and then positioned *∼*5 mm and 4 mm apart from each other on the axial and lateral planes, respectively. Then a solution mixture of agar, intralipid, and silica beads with a respective weight ratio of 1.5%, 1%, and 1% was poured into the tank to provide the ultrasound background speckles as well as the optical scattering [65]. While ultrasound imaging contrast of the phantom was based on the acoustic impedance differences between the wire and the background, the photoacoustic imaging contrast was based on the light absorption of metal wire targets dyed with India ink.

### 4.8 Field-of-view targets

The FOV of PA imaging was characterized by imaging India ink dyed micro wire targets placed orthogonally to the imaging plane of an area of 20 mm wide by 25 mm deep. Micro metal wires were spaced 5 mm and 3 mm between each other in the axial and lateral planes, respectively, to minimize interference of PA signal while maintaining the sufficient density of the targets. Then the FOV characterization was performed in an optical scattering medium made of deionized water with 1% intralipid. PA imaging was conducted at 750 nm wavelength.

### 4.9 Tube phantom of flowing blood and static ICG

To validate the functional and molecular imaging capabilities of UFUS and MSPA modalities of the fUSPA system, we developed a tube phantom consisting of two polyethylene tubes (outer diameter: 2.08 mm, inner diameter: 1.57 mm, PE 205, BD Medical, Franklin Lakes, NJ, USA). One tube (left side in Fig. S5a) was filled with defibrinated bovine blood (Lampire biological laboratories, Pipersville, PA, USA) while circulated using a peristatic pump (model 3386, Cole-Parmer, Vernon Hills, IL, USA). The other tube (right side in Fig. S5a) was filled with 1290 *µ*M ICG in deionized water. The tubes were embedded inside a solid phantom, made of 1.5% agar and 1% intralipid to mimic optical scattering, to reduce any motion artifacts caused by the pump. To induce the blood oxygenation differences, a gas chamber was added to the blood circulation pathway in the left tube. The gas chamber was filled with a mixture of CO_2_ and O_2_ with their respective ratio controlled by individual flow meters. The output flow rate was maintained constant at 3 L/min. For example, 100% O_2_ was achieved by giving O_2_ at a flow rate of 3 L/min, while a mixture of 66% O_2_ and 33% CO_2_ was achieved by giving O_2_ at 2 L/min and CO_2_ at 1 L/min. As previously described in methods 4.2, the interleaved MSPA (with 750 nm, 800 nm, and 850 nm) and UFUS data acquisition of the tube phantom was performed using the fUSPA imaging system. While the MSPA data of each gas mixture concentration was processed with the methods described in 4.3 to provide unmixed maps of HbO, HbD, SO2, and ICG molecular distribution, the UFUS data was processed with methods described in 4.4 to generate the blood flow map.

### 4.10 Animal handling and surgery

All animal procedures were approved by the Pennsylvania State University Institutional Animal Care and Use Committee (IACUC). Adult Sprague Dawley rats (n=4, 10 - 16 weeks old) were used in the experiments and acclimated for at least 5 days prior to the surgery. These animals received craniotomy survival surgery prior to imaging sessions to allow US and PA imaging of the brain. On the date of the surgery, the animal was anesthetized by administering isoflurane/O_2_ via vaporizer at 3-5% and maintained at 1-2% throughout the surgery.

A sagittal skin incision was performed across the posterior part of the head to expose the skull, and a dental drill was used to mark the window for craniotomy with a size up to 10 mm *×* 14 mm. Beginning with a 1.0 mm drill bit, a rectangular cut was made within the marked borders of the craniotomy. The drilling was frequently paused for cooling the skull bone with cold saline to reduce heat damage, edema, and to control bleeding around or underneath the skull. The thinning was continued until the pial vessels were clearly seen through the thinned skull, the bone flap was lifted off using fine tip forceps and exposed the underlying intact dura. Then 1.5% weight ratio agar (UltraPure Agarose, Invitrogen, Waltham, MA, USA), mixed in sterilized saline was molded into a cuboid with a similar thickness to the skull and cut into a size slightly smaller than the window. The agar cuboid was then placed on top of the dura to provide hydraulic pressure and acoustic coupling. Then the whole window on the animal head was covered by a 50 *µ*m thick PMP film (Goodfellow, Pittsburgh, PA, USA) as a skull prosthetic and acoustic transparent window. Next, the acoustic window was sealed to the surrounding skull using dental cement (C&B metabond, Parkell Inc., Brentwood, NY, USA). A nut was placed at the Bregma as an anatomical marker for in vivo imaging sessions. Following the surgery, the animal was allowed one week to recover before the start of any imaging experiments.

### 4.11 Animal imaging and CO**_2_** stimulation

At the beginning of the imaging session, the animal was anesthetized using 3% isoflurane mixed with medical oxygen at 2 L/min. Once the animal was stably anesthetized, it was moved to the imaging stage and fixed to the stereotaxic frame with a nose cone providing a continuous supply of 1 - 1.5% isoflurane mixed with medical oxygen at 2 L/min to maintain anesthesia throughout the imaging session. Then the imaging window was gently cleaned with sterilized saline using cotton-tipped applicators. Warm ultrasound coupling gel mixed with saline was then applied over the window to couple the fUSPA probe to conduct UFUS, MSPA and ULM imaging. During the UFUS and MSPA imaging sessions, we performed the hypercapnia (CO_2_) by delivering 5% CO_2_ mixed with oxygen through the nose cone. The 5% CO_2_ volume fraction was controlled by adjusting the flow rate of the O_2_ and CO_2_ using individual flow meters. UFUS and MSPA imaging data was continuously acquired during an 8-minute stimulation paradigm, which is composed of a 2-minute baseline, a 3-minute CO_2_ stimulation period, and a 3-minute post-stimulation period.

For ULM imaging, a 26 Gage intravenous (IV) catheter needle was placed in the rat tail vein for MB injection. MBs (SIMB 4-5, Advanced microbubbles, Newark, CA, USA) were diluted to 10^8^ MBs/mL following the manufacturer’s protocol. Two bolus injections of 400 *µ*L MBs were performed at t = 0 (at the start of ULM imaging) and t = 5 minutes, respectively. Then, as described above, the ULM imaging sequence was performed to obtain super-resolved microvasculature images of the brain.

To investigate the exogenous photoacoustic molecular imaging capabilities of the fUSPA imaging system, Food and Drug Administration (FDA) approved optical fluorescence contrast agent, ICG, was intravenously administered while the animal was maintained under the isoflurane anesthesia. The 10-minute MSPA imaging protocol was performed to delineate the depth dependent intrinsic (HbO and HbD) and exogenous (ICG) molecular contrast information. After a 1-minute baseline acquisition of MSPA imaging data, a 500 *µ*L of 129 *µ*M ICG in sterilized saline was slowly injected through the catheterized tail vein (*∼* 16.7 *µ*L/second). Following the completion of the injection, MSPA imaging sequence was continuously performed for 8:30 minutes to study post-injection time activity of intravascular ICG inside the brain.

### 4.12 Offline processing for continuous UFUS and MSPA data

The saved UFUS and MSPA data were processed offline to extract the time dynamics of the rCBV, rSO2 (or relative ICG changes, e.g. Fig. 3e), and CBF changes. While UFUS data were processed to obtain the CBV and CBF maps of each time point, MSPA data were processed to obtain the SO2 map of each time point. Relative change in the intensity of CBV maps of each pixel is calculated by:

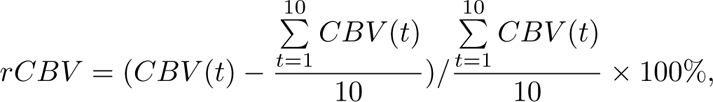

where CBV(t) denotes the power Doppler intensity or CBV intensity of the pixel at time t obtained from the power Doppler intensity. Similarly, rCBF, and rSO2 were calculated by:

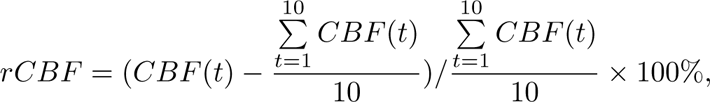

and

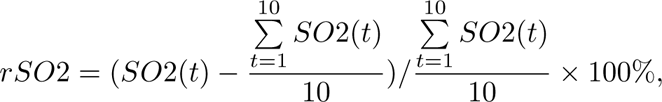

respectively. The above calculation was done for each pixel to generate relative CBV, CBF, and SO2 change maps and videos.

### 4.13 Ex vivo isolated rat brain imaging

We performed ex vivo whole rat brain imaging experiments to help understand and optimize fUSPA device imaging capabilities for in vivo rat brain imaging. The rat brain was carefully removed from the head following CO_2_ euthanasia according to the approved IACUC protocol. Immediately after the removal, the whole brain was embedded inside a 1.5% agar medium to provide additional stability and easy coupling to the fUSPA device. Next, dual-modality US and MSPA imaging of the ex vivo brain was performed using the fUSPA system.

### 4.14 Cross-correlation function for cortex-vein temporal dynamics

To investigate the coupling between the temporal dynamics of the veins and the overall cortical vessels, we performed continuous UFUS imaging of the rat brain at high temporal resolution (*∼* 0.71 seconds per frame). The animal was maintained under anesthesia using 1 - 1.5% isoflurane while the continuous UFUS imaging was performed for 5 minutes without any stimulation. The acquired data were then analyzed offline to investigate the correlation between the venous and cortical vessel CBV activities. As shown in Fig. 5, we marked four ROIs for further analysis. The ROI surrounding the cortex is denoted as ROI 1, and ROI surrounding a single vein (as confirmed with ULM, Fig. 5c) is denoted as ROI 2. ROI 3 in the cortex encompasses ROI 2 and was used to help visualize the anti-correlation effect from the vein shown in ROI 2 (Fig. 5g). ROI 4 marks the ventricle region. The temporal dynamics of the cortex and vein were obtained by averaging the time activities of the ROI 1 and ROI 2 voxels, respectively. The example averaged time series are shown in Fig. 5d. The cross-correlation function was done for two scenarios: 1) cortex-all correlation, where the temporal dynamics of each individual voxel was cross correlated with the time averaged cortical dynamics (from ROI 1), and the resulted correlation coefficient (R) was plotted as the map for different lags (Fig. 5e and Fig. S13). 2) cortex-vein correlation, where the cross-correlation was calculated between the averaged cortical vascular dynamics from ROI 1 and the averaged venous dynamics from ROI 2 (Fig. 5f-g).

## Acknowledgments

We thank Eugene Gerber for his kind support in machining and Vaishnavi Muralidharan for her support in manuscript proofreading and reviewing. We also acknowledge funding from various sources, including the Penn State Cancer Institute seed grant (S.-R.K.), and NIH cross disciplinary neural engineering training program T32NS115667 (B.J.G., H.C.).

## Author Contributions

Conceptualization, H.C., and S.-R.K.; software and data analysis, H.C., S.M., P.G., S.A., X.L., and T.X.; investigation, X.L., P.J.D., N.Z., B.J.G., and S.-R.K.; data curation, H.C., S.M., and V.N.; writing—original-draft preparation, H.C., S.M.; experiment design, H.C., Q.L., and S.-R.K.; hardware, H.C., M.L., J.L., B.J.G., and W.T.; supervision, S.-R.K.; project administration, S.-R.K.; funding acquisition, B.J.G., and S.-R.K

## Data Availability

Data underlying the results presented in this paper are not publicly available at this time but may be obtained from the authors upon reasonable request.

