## Supplementary Materials for "Dissecting Multiparametric Cerebral Hemodynamics using Integrated Ultrafast Ultrasound and Multispectral Photoacoustic Imaging"

### Supplementary Figures

**Fig. S1: The schematics and photograph of fUSPA described in Fig. 1.**

**Fig. S2: Timing diagram of acquisition sequences for interleaved ultrafast ultrasound (UFUS) and multispectral photoacoustic (MSPA) imaging using fUSPA.**

**Fig. S3: Ultrasound (US) pulse-echo and photoacoustic (PA) signal characterization of the 128-element ultrasound transducer using a flat metal target placed inside water medium.**

**Fig. S4: Ultrasound (US) and photoacoustic (PA) imaging characterization of fUSPA using microwire targets.**

**Fig. S5: Ultrasound (US), multispectral photoacoustic (MSPA), and ultrafast ultrasound (UFUS) based power Doppler imaging validation using 2.08 mm diameter silicon tubes containing flowing blood and static indocyanine green (ICG).**

**Fig. S6: Multispectral photoacoustic (MSPA) images of the three coronal brain imaging locations shown in Fig. 2.**

**Fig. S7: Ultrasound (US) and photoacoustic (PA) imaging of isolated ex vivo rat brain.**

**Fig. S8: Multispectral photoacoustic (MSPA) images of the pre- and post-intravenous injection of indocyanine green (ICG), corresponding to Fig. 3.**

**Fig. S9: Single vessel relative cerebral blood volume (rCBV) and relative cerebral blood flow (rCBF) analysis, corresponding to the region-of-interest in Fig. 4g, reveals functional differences between arterioles and venules during hypercapnia stimulation.**

**Fig. S10: fUSPA imaging revealed multiparametric hemodynamic changes at the superior sagittal sinus during hypercapnia stimulation.**

**Fig. S11: fUSPA imaging revealed multiparametric hemodynamic changes at the ventricle region during hypercapnia stimulation.**

**Fig. S12: Ultrasound (US) and photoacoustic (PA) frames of the sagittal brain section near the sagittal midline corresponding to Fig. 5**

**Fig. S13: The maps of cross-correlation between the averaged temporal dynamics of the cortex (ROI 1 in Fig. 5) and the temporal dynamics of each voxel in the image, corresponding to Fig. 5.**

**Fig. S14: The schematic for the laser light refraction through the custom designed acrylic lens and associated focal distance calculations for the fUSPA imaging device.**

#### Supplementary Notes

**Supplementary Note 1: Light refraction and focal distance calculation for the fUSPA imaging device**

**Supplementary Note 2: fUSPA system characterization and validation**

#### Supplementary Movies

**Movie S1: Ultrafast ultrasound based cerebral blood volume (CBV) imaging of rat brain and relative CBV change map during hypercapnia stimulation, corresponding to Fig. 4.**

**Movie S2: Ultrafast ultrasound based descending and ascending cerebral blood flow maps in brain during hypercapnia stimulation, corresponding to Fig. 4.**

**Movie S3: Multispectral photoacoustic derived oxygen saturation and relative oxygen saturation change map during hypercapnia stimulation, corresponding to Fig. 4.**

#### Supplementary Figures

**Fig. S1**

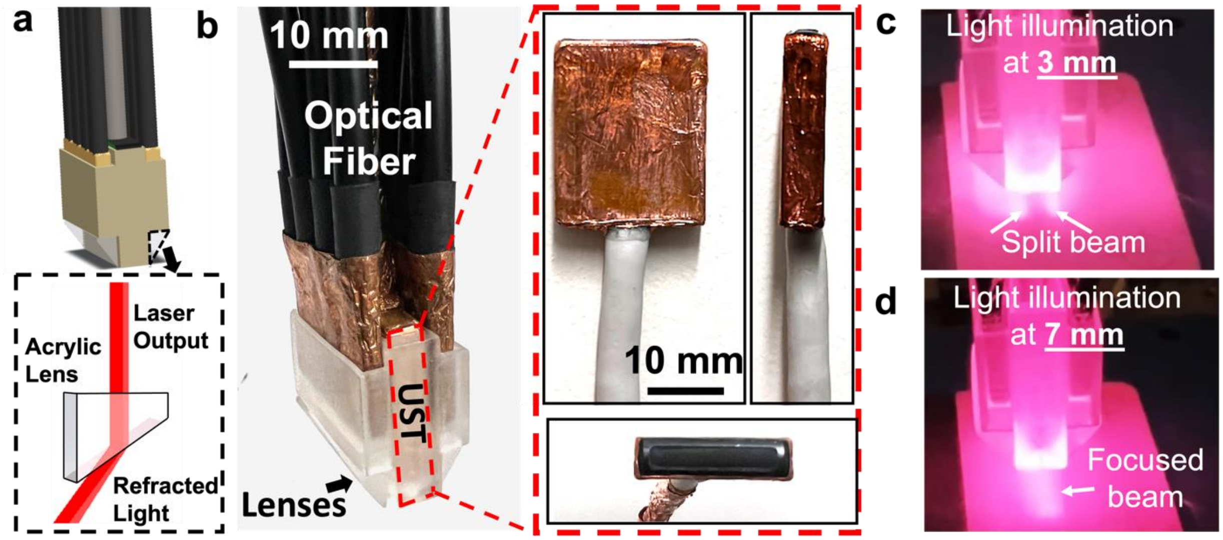


**Fig. S1: The schematics and photograph of fUSPA described in Fig. 1. (a)** The schematic of the 3D-printed holder that integrated the high-frequency ultrasound transducer (UST) probe and optical fibers. The acrylic lenses at the bottom of the holder (bottom Fig. S1a) refract the fiber optic beam to the ultrasound detection zone. **(b)** The photograph of the integrated fUSPA imaging device. The red inlet shows the zoomed in pictures of the high frequency UST. **(c-d)** Pictures demonstrate the optical beam profile at ~3 mm and ~7 mm distance below the imaging device.

**Fig. S2**

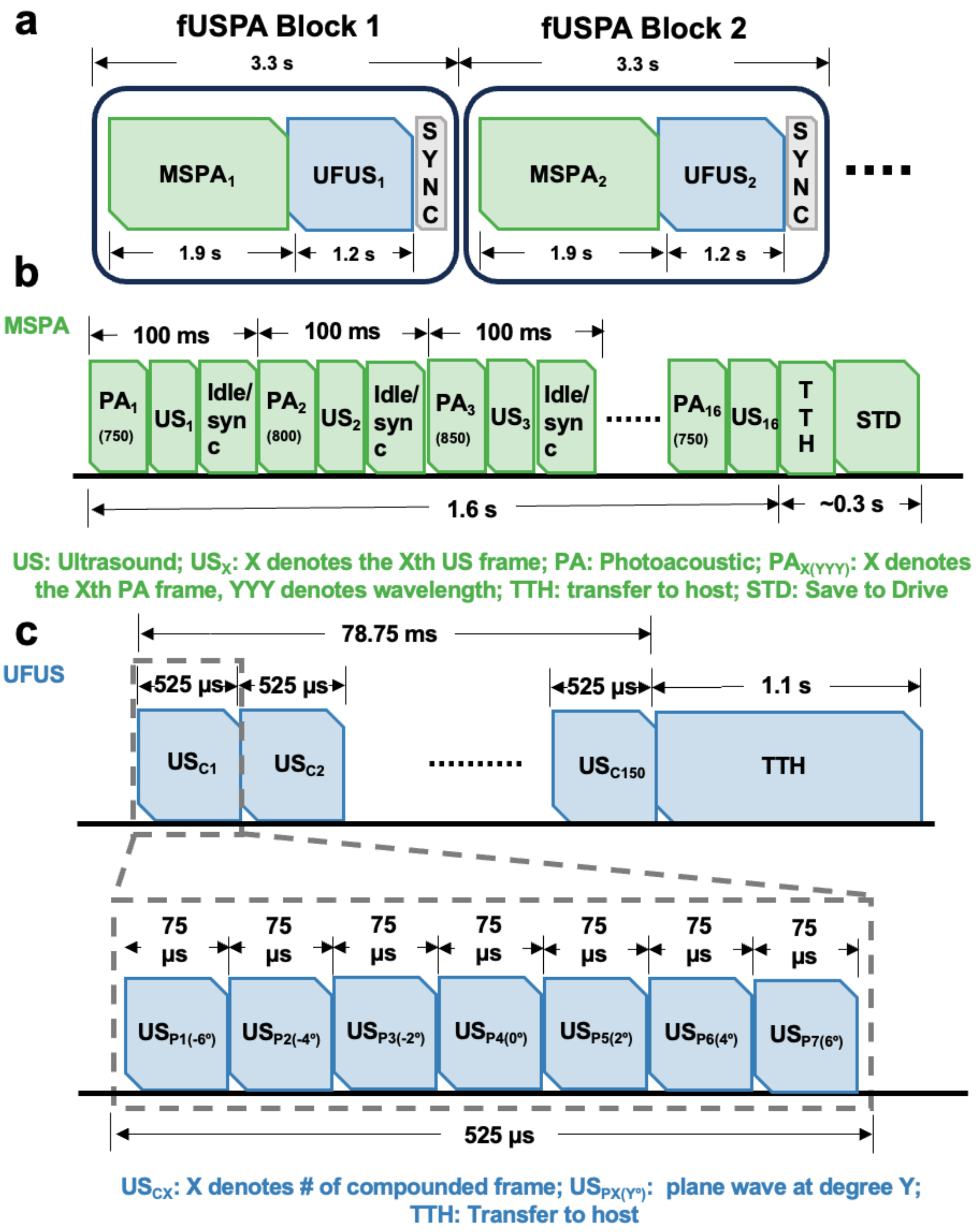


**Fig. S2: Timing diagram of acquisition sequences for interleaved ultrafast ultrasound (UFUS) and multispectral photoacoustic (MSPA) imaging using fUSPA.** **(a)** Each fUSPA block sequence was consisted of a ~1.9 s MSPA acquisition and a ~1.2 s UFUS acquisition, followed by ~0.2 s of SYNC which triggers the acquisition of the next consecutive fUSPA block. The resultant fUSPA frame rate was ~3.3 seconds per frame. **(b)** Each MSPA block consisted of three PA data acquisitions at three wavelengths (750 nm, 800 nm, and 850 nm) for linear spectral unmixing. This sequence was repeated five times, resulting in a total of 15 PA frames (5 frames per each of 3 wavelengths). One additional PA frame was acquired (and discarded) at the beginning to improve synchronization of laser firing and subsequent PA data acquisition. Therefore, with 10Hz laser pulse repetition frequency, the MSPA data acquisition took 1.6 s for 16 PA frames and 0.3 s to transfer the data from Verasonics to the local solid-state drive. **(c)** For the UFUS sequence, 150 compounded US images were acquired to generate one CBV or CBF map. Each compounded image was formed by transmitting and receiving 7 angled plane waves (-6°, -4°, -2°, 0°, 2°, 4°, 6°) with 75 μs between each angle, which resulted in a frame rate of 525 μs per compounded US frame. Therefore for 150 compounded US frames, one UFUS based image acquisition took 78.75 ms with a transfer to host (TTH) time of ~1.1 s.

**Fig. S3**

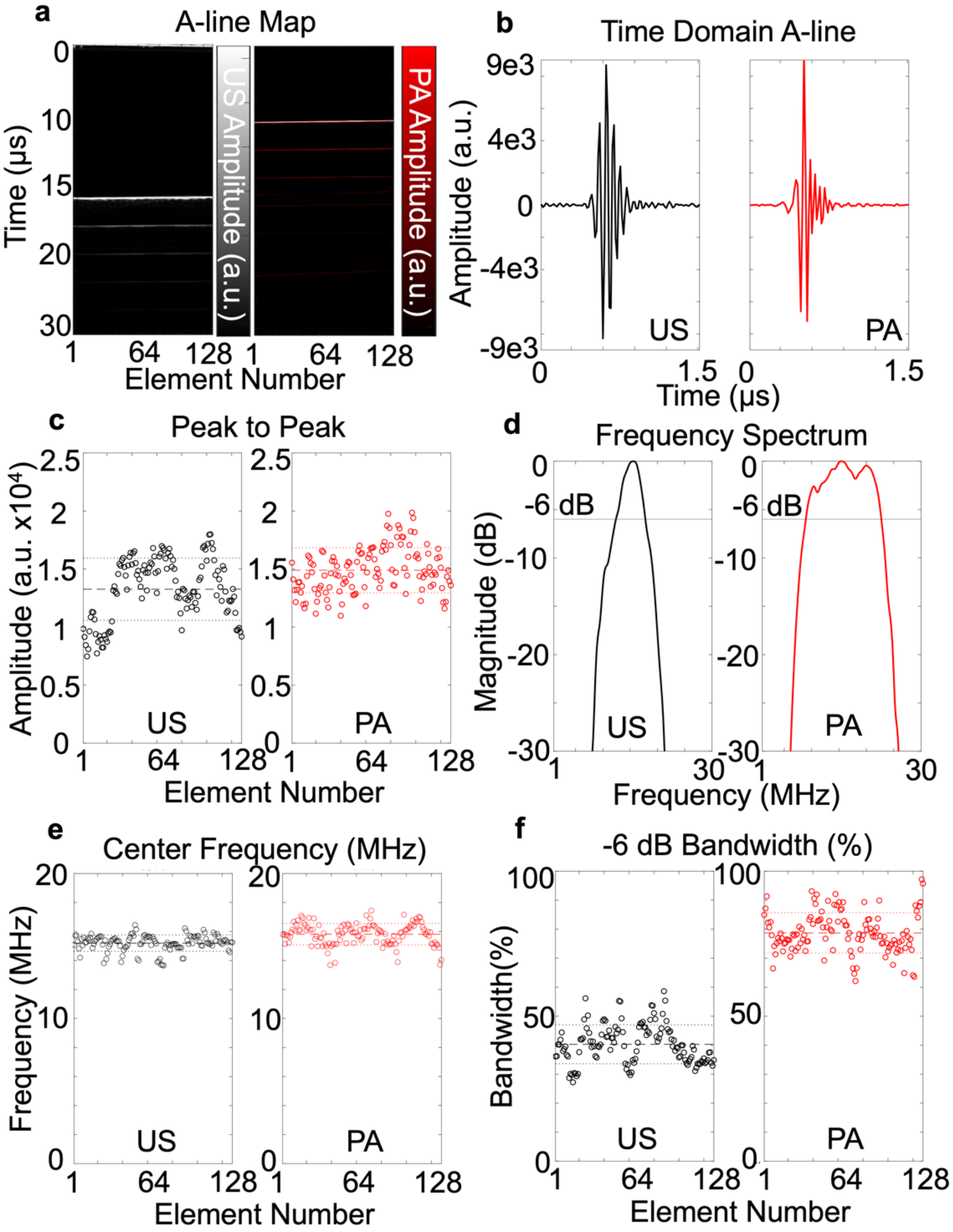


**Fig. S3: Ultrasound (US) pulse-echo and photoacoustic (PA) signal characterization of the 128-element ultrasound transducer using a flat metal target placed inside water medium. (a)** A-line maps of the US and PA signals received from the metal target shows near flat response for all elements. The double the time taken for the US A-line is due to round trip (transducer to target and target to transducer) of US pulse. **(b)** The example time-domain US and PA A-line signals from the center element #64 of the transducer. **(c)** Scattered points represent the US and PA A-line peak-to-peak amplitude corresponding to each of the 128 transducer elements. The center dashed lines in US and PA plots denote the averaged amplitude, with top and bottom dashed line denote the average amplitude +/- standard deviation. **(d)** The frequency responses of the

US and PA A-lines, obtained from Fourier transform of the respective signals in (b), show the respective center frequencies and the -6 dB bandwidths. **(e-f)** The dot plot of the measured center frequencies and the -6 dB bandwidths of each element from US pulse-echo and PA receiving mode, respectively. The center dashed lines denote the averaged frequency/bandwidth, with the top and bottom dashed lines denote the average center frequency/bandwidth +/- standard deviation.

**Fig. S4**

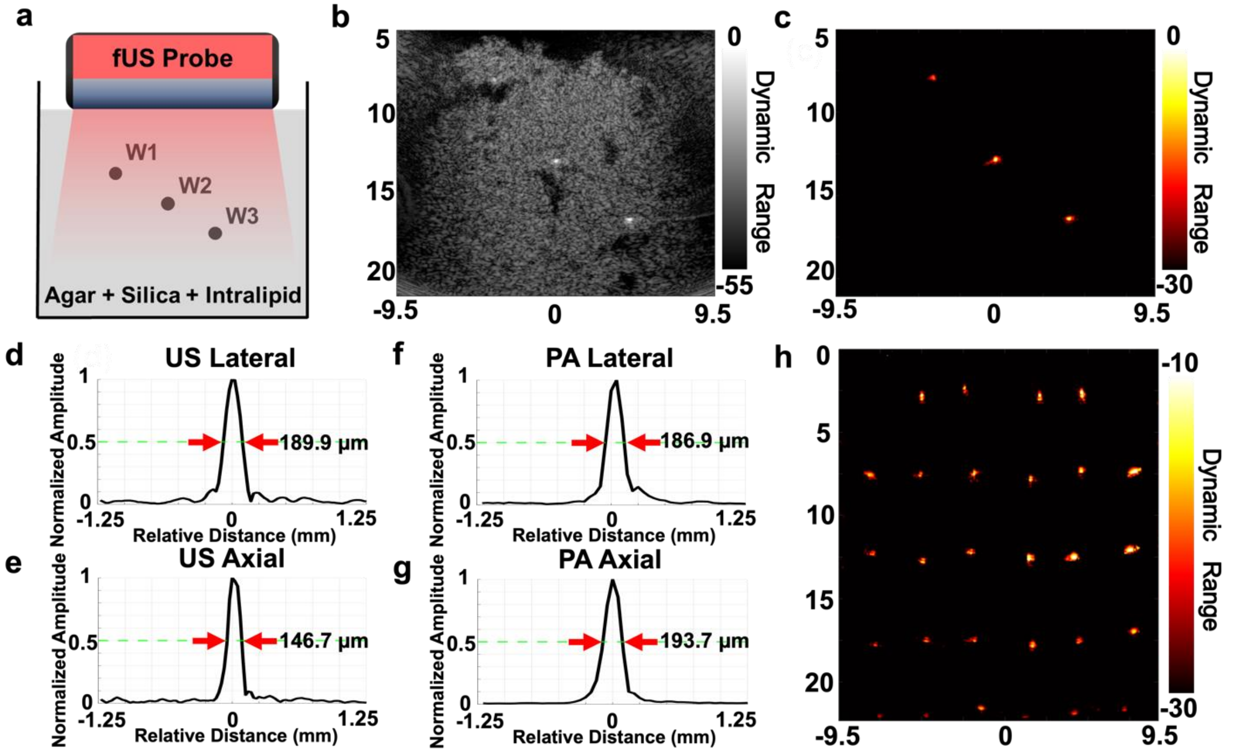


**Fig. S4: Ultrasound (US) and photoacoustic (PA) imaging characterization of fUSPA using microwire targets. (a)** The schematic of the resolution phantom with three 50 μm diameter microwires (W1-W3) embedded inside a tissue mimicking scattering medium consisted of 1.5% agar, 1% silica particles (for acoustic scattering) and 1% intralipid (for optical scattering). **(b)** The US image and **(c)** the PA image of the resolution phantom. **(d-g)** To characterize imaging resolution, the full-width-half-maxima (FWHM) of the center wire target in US and PA images were calculated in both axial and lateral directions. **(h)** The microwire targets were distributed uniformly in a 25 mm deep by 20 mm wide area to characterize the field of view (FOV).

**Fig. S5**

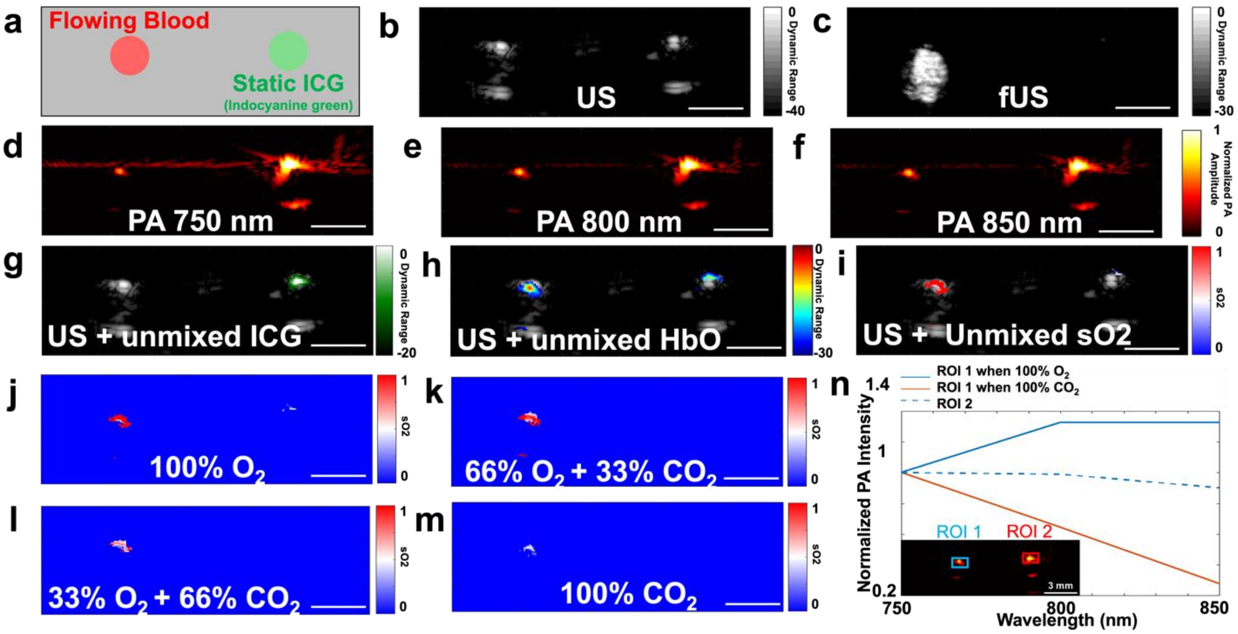


**Fig. S5: Ultrasound (US), multispectral photoacoustic (MSPA), and ultrafast ultrasound (UFUS) based power Doppler imaging validation using 2.08 mm diameter silicon tubes containing flowing blood and static indocyanine green (ICG). (a)** A schematic view of the phantom with left tube containing flowing bovine blood mixed with a controlled gas mixture and right side with a tube containing static ICG at 1290 µM concentration. **(b)** A cross-sectional (B-mode) US image of the tube phantom showed anatomical contrast generated from both tubes due to acoustic impedance mismatch. **(c)** Power Doppler image showed the blood flow in the left tube due to flowing particles. MSPA images of the phantom acquired at three wavelengths **(d)** 750 nm, **(e)** 800 nm, and **(f)** 850 nm. ICG, HbO, and HbD molecular information was unmixed using the three-wavelength MSPA images and co-registered with US image to generate **(g)** US + ICG, **(h)** US + HbO, and **(i)** US + SO2 images. By varying the gas mixtures delivered to the flowing bovine blood, the unmixed MSPA images of the SO2 showed a descending oxygen saturation corresponding to the reduced oxygen and increased carbon dioxide level: **(j)** 100% $O_2$, **(k)** 66% $O_2$ + 33% $CO_2$, **(l)** 33% $O_2$ + 66% $CO_2$, and **(m)** 100% $CO_2$. Color bar indicates normalized SO2 values from 0 to 1. **(n)** The quantitative comparison of normalized PA intensity vs. wavelength in the blood tube (ROI 1) under 100% $O_2$ (blue solid line) and 100% $CO_2$ (blue dashed line). ROI 2 shows the normalized PA intensity vs. wavelength in the 1290 µM ICG tube (red solid line). HbO: oxygenated hemoglobin; HbD: deoxygenated hemoglobin; SO2: oxygen saturation; ROI: region-of-interest. Scale bars represent 2 mm.

**Fig. S6**

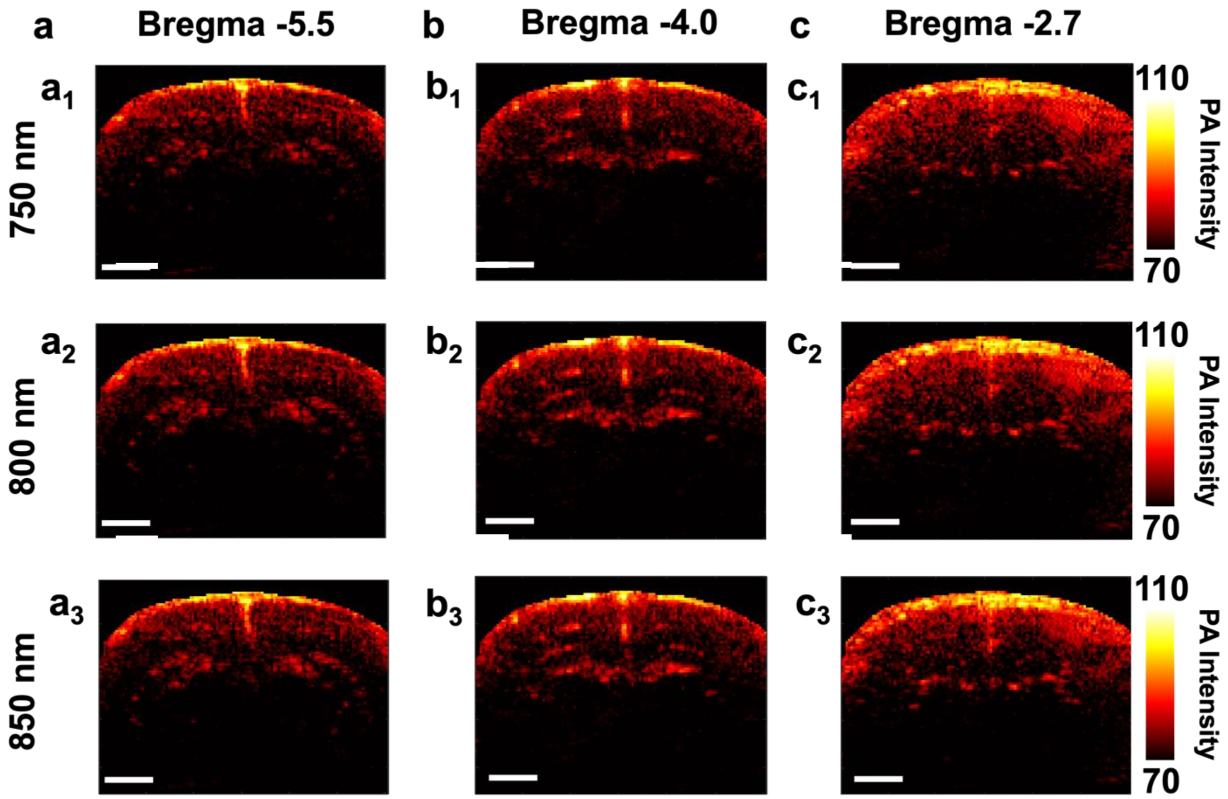


**Fig. S6: Multispectral photoacoustic (MSPA) images of the three coronal brain imaging locations shown in Fig. 2. (a)** The **($a_1$)** 750 nm, **($a_2$)** 800 nm, and **($a_3$)** 850 nm PA images of the Bregma -5.5 location. **(b)** The **($b_1$)** 750 nm, **($b_2$)** 800 nm, and **($b_3$)** 850 nm PA images of the Bregma -4.0 location. **(c)** The **($c_1$)** 750 nm, **($c_2$)** 800 nm, and **($c_3$)** 850 nm PA images of the Bregma -2.7 location. Scale bars represent 2 mm.

**Fig. S7**

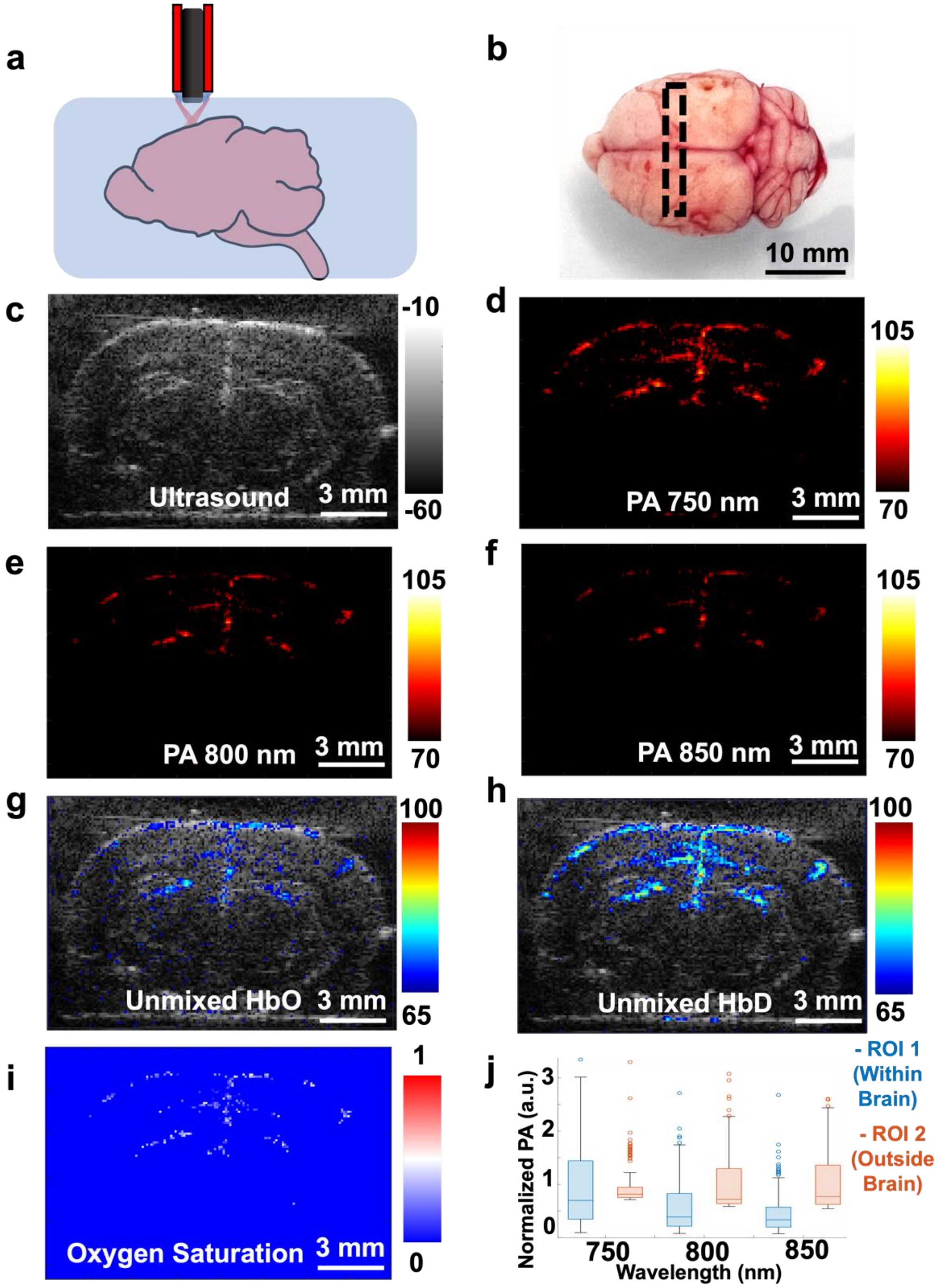


**Fig. S7: Ultrasound (US) and photoacoustic (PA) imaging of isolated ex vivo rat brain. (a)** A schematic view of the agar phantom embedded with the whole rat brain. The fUSPA imaging head is coupled to the phantom using ultrasound gel. **(b)** Picture of the whole rat brain with dotted rectangular box showing the orientation of the fUSPA system to generate images along the coronal plane**. (c)** B-mode US image of the brain and corresponding MSPA images at **(d)** 750 nm, **(e)** 800 nm, and **(f)** 850 nm wavelengths. **(g)** HbO and **(h)** HbD maps, obtained from the linear spectral unmixing of MSPA images, were co-registered with the grayscale US image**. (i)** The oxygen saturation map was generated from the unmixed HbO and HbD images by calculating the pixel-value intensity ratio HbO/(HbO + HbD). **(j)** Plots of PA intensities as function of wavelength for ROIs inside (ROI 1) and outside (ROI 2) the brain, showed clear differences with ROI 1 following a typical HbD optical absorption characteristic curve while ROI 2 having similar intensities across three wavelengths. MSPA: multispectral photoacoustic; HbO: oxygenated hemoglobin; HbD: deoxygenated hemoglobin; ROI: region-of-interest.

**Fig. S8**

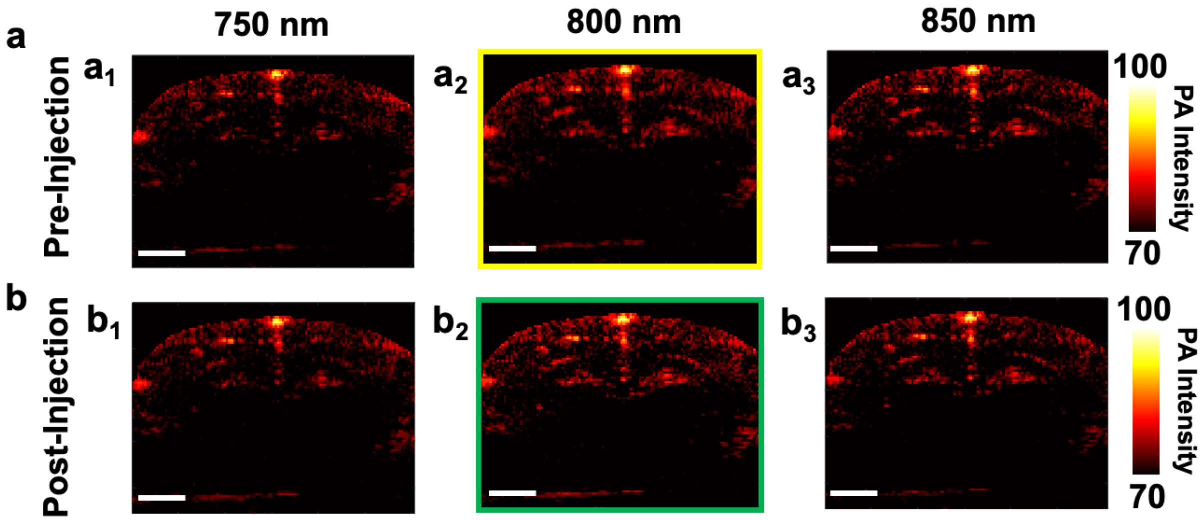


**Fig. S8: Multispectral photoacoustic (MSPA) images of the pre- and post-intravenous injection of indocyanine green (ICG), corresponding to Fig. 3. (a)** The PA images acquired at **($a_1$)** 750 nm, **($a_2$)** 800 nm, and **($a_3$)** 850 nm wavelengths prior to ICG injection, and **(b)** the **($b_1$)** 750 nm, **($b_2$)** 800 nm, and **($b_3$)** 850 nm PA images of the same imaging location after ICG injection**.** The 800 nm post-injection PA frame, green outlined ($b_2$), show increased PA intensity compared to pre-injection 800 nm PA image shown in yellow outlined ($a_2$). Scale bars represent 2 mm.

**Fig. S9**

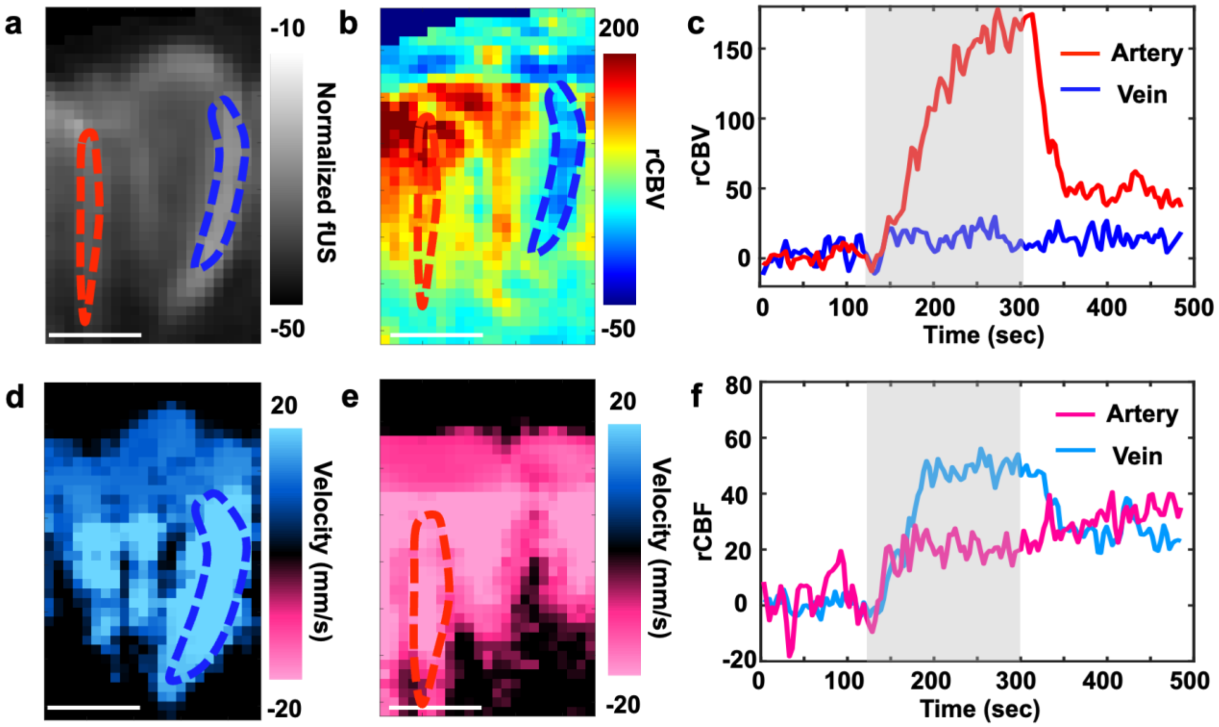


**Fig. S9: Single vessel relative cerebral blood volume (rCBV) and relative cerebral blood flow (rCBF) analysis, corresponding to the region-of-interest in Fig. 4g, reveals functional differences between arterioles and venules during hypercapnia stimulation.** **(a)** The UFUS based CBV image of the ROI, with blue and red dashed lines marking the venule and the arteriole respectively. **(b)** The rCBV change in the ROI during hypercapnia (t = 260 second). **(c)** The plots of averaged rCBV change during hypercapnia stimulation experiment with gray shaded area indicating $CO_2$ ON period of 180 seconds (from 120 seconds to 300 seconds) for the delineated venule and arteriole. **(d)** UFUS based ascending CBF map and **(e)** UFUS based descending CBF map. **(f)** The rCBF change obtained from the single vessel delineated in blue and red dashed lines in (e) and (f). UFUS: ultrafast ultrasound; CBV: cerebral blood volume; rCBV: relative cerebral blood volume; ROI: region-of-interest; CBF: cerebral blood flow; rCBF: relative blood flow. Scale bars represent 0.5 mm.

**Fig. S10**

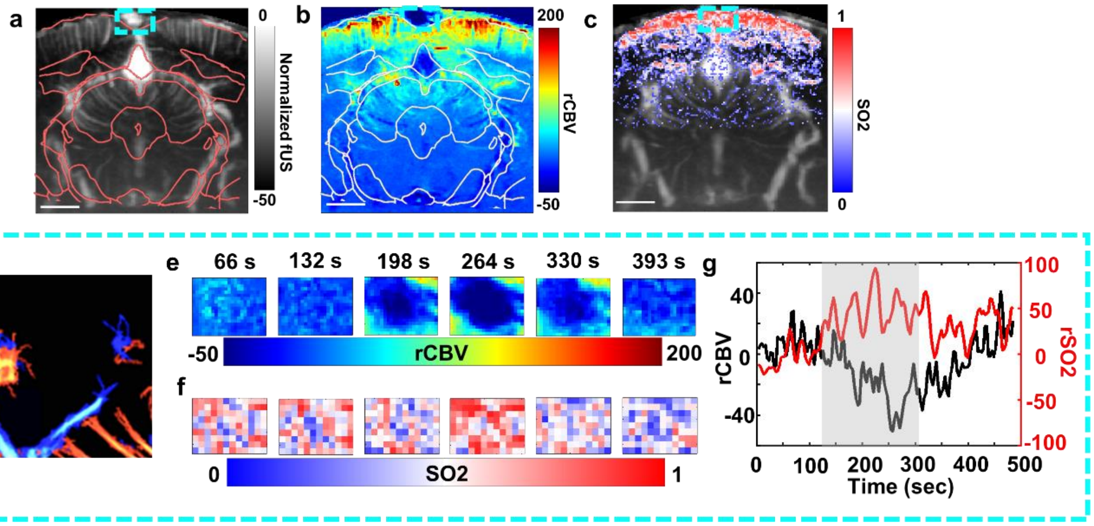


**Fig. S10: fUSPA imaging revealed multiparametric hemodynamic changes at the superior sagittal sinus during hypercapnia stimulation. (a)** The grayscale UFUS based CBV image overlaid with the atlas at Bregma -5.8. **(b)** The rCBV change in the ROI during hypercapnia (t = 260 second). **(c)** The SO2 map, obtained from the spectral unmixing of MSPA imaging data, is overlaid onto the corresponding CBV map. The cyan dashed box in (a-c) delineates the ROI for the superior sagittal sinus. The scale bars represent 2 mm. **(d)** The enlarged ULM at the corresponding ROI**. (e)** The enlarged rCBV change map and **(f)** the enlarged unmixed SO2 map at the given ROI for different time points during the hypercapnia stimulation paradigm. **(g)** The plots of averaged rCBV changes (black line) and the averaged rSO2 changes (red line) in the given ROI during hypercapnia stimulation paradigm, with gray shaded area indicating $CO_2$ ON period of 180 seconds (from 120 seconds to 300 seconds). CBV: cerebral blood volume; rCBV: relative cerebral blood volume; ROI: region-of-interest; MSPA: multispectral photoacoustic; SO2: oxygen saturation; rSO2: relative oxygen saturation.

**Fig. S11**

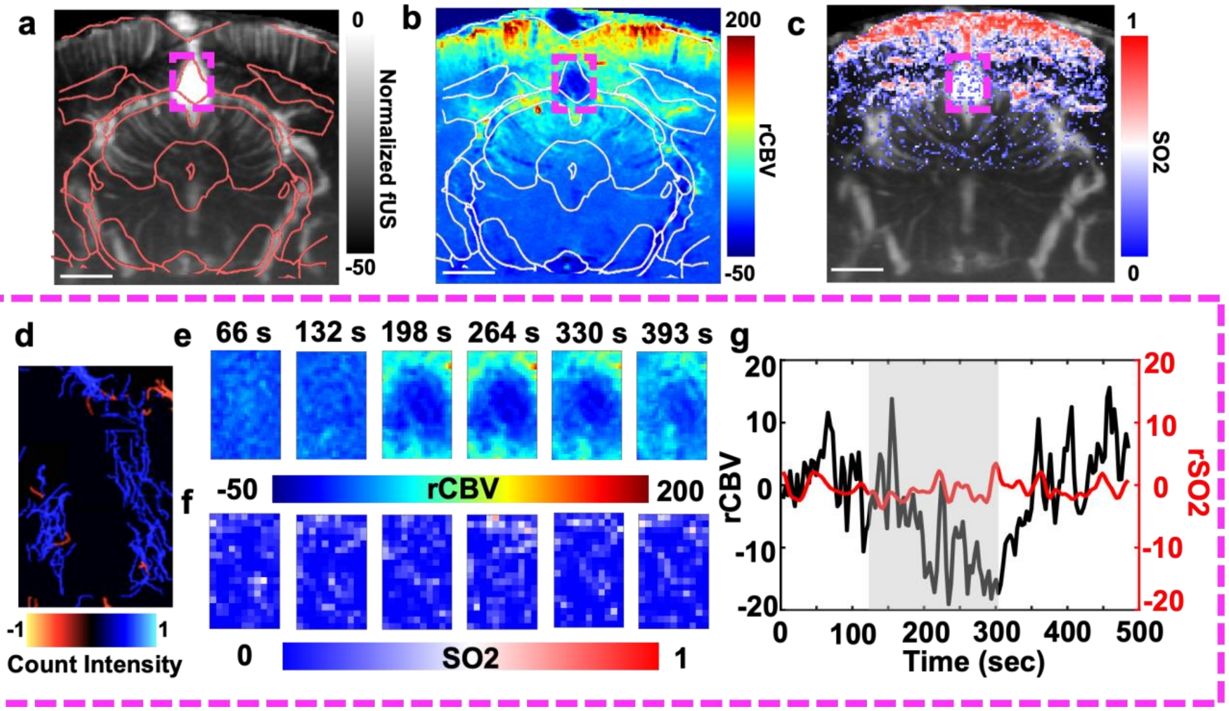


**Fig. S11: fUSPA imaging revealed multiparametric hemodynamic changes at the ventricle region during hypercapnia stimulation. (a)** The grayscale UFUS based CBV image overlaid with the atlas at Bregma -5.8. **(b)** The rCBV change in the ROI during hypercapnia (t = 260 second). **(c)** The SO2 map, obtained from the spectral unmixing of MSPA imaging data, is overlaid onto the corresponding CBV map. The magenta dashed box in (a-c) delineates the ROI for the ventricle region. The scale bars represent 2 mm. **(d)** The enlarged ULM at the corresponding ROI**.** **(e)** The enlarged rCBV change map and **(f)** the enlarged unmixed SO2 map at the given ROI for different time points during the hypercapnia stimulation paradigm. **(g)** The plots of averaged rCBV changes (black line) and the averaged rSO2 changes (red line) in the given ROI during hypercapnia stimulation paradigm, with gray shaded area indicating $CO_2$ ON period of 180 seconds (from 120 seconds to 300 seconds). CBV: cerebral blood volume; rCBV: relative cerebral blood volume; ROI: region-of-interest; MSPA: multispectral photoacoustic; SO2: oxygen saturation; rSO2: relative oxygen saturation.

**Fig. S12**

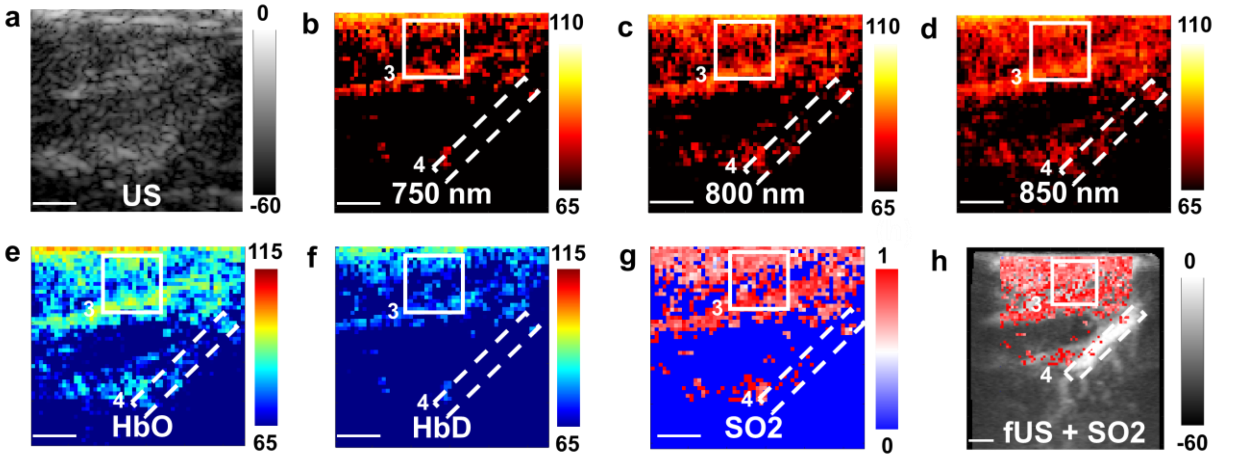


**Fig. S12: Ultrasound (US) and photoacoustic (PA) frames of the sagittal brain section near the sagittal midline corresponding to Fig. 5. (a)** B-mode US image of the sagittal brain region. **(b-d)** MSPA frames at the same location imaged with 750 nm, 800 nm, and 850 nm wavelengths respectively. The three wavelengths were linearly unmixed to generate **(e)** the HbO map and **(f)** the HbD map. By calculating the ratio of HbO and HbD map, **(g)** SO2 map was generated, **(h)** and the SO2 map was further co-registered with the corresponding power Doppler based CBV map. ROI 3 covers a cortical brain region that is also shown in Fig.5, and ROI 4 delineates the ventricle region HbO: oxygenated hemoglobin; HbD: deoxygenated hemoglobin; SO2: oxygen saturation; ROI: region-of-interest. Scale bars represent 1 mm.

**Fig. S13**

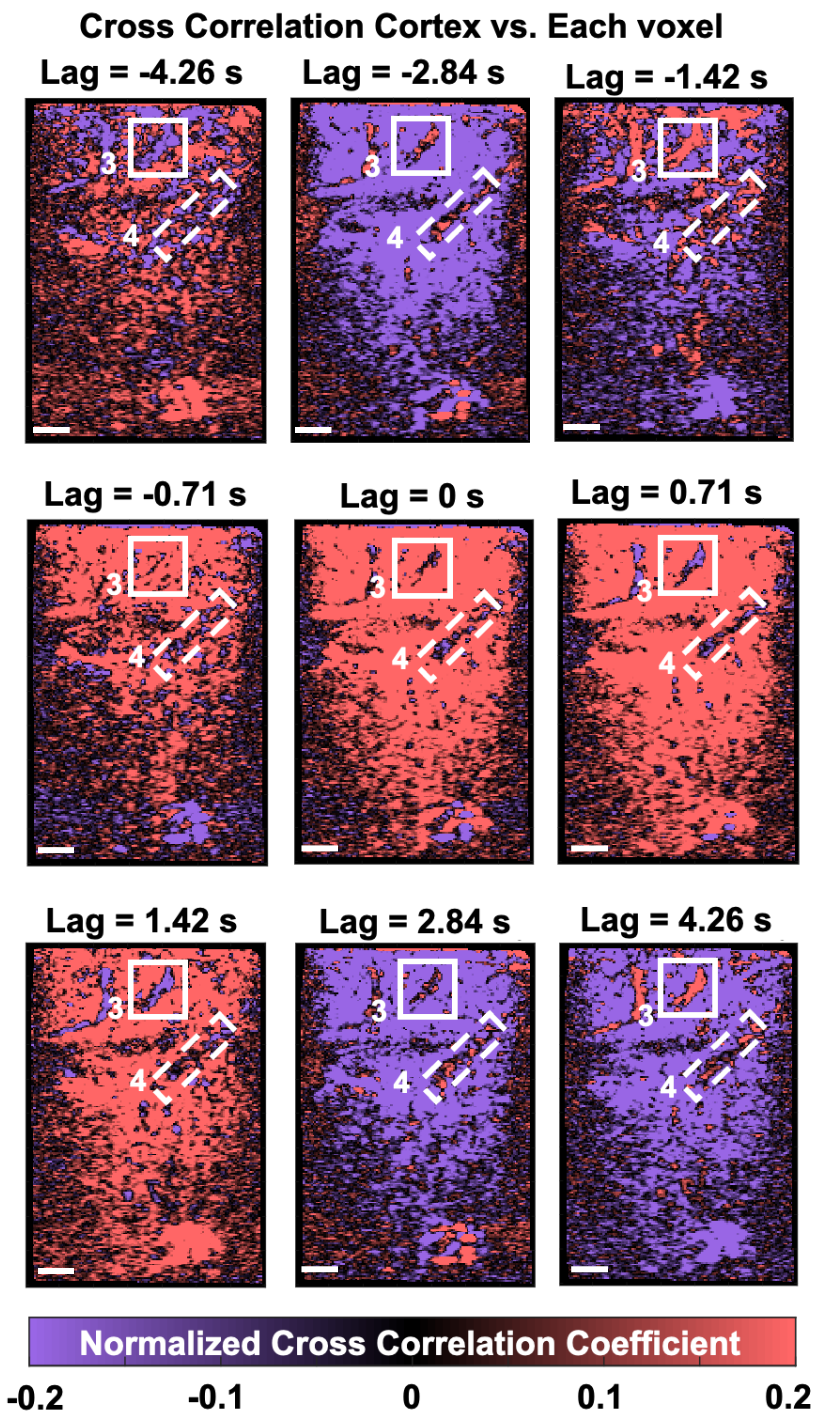


**Fig. S13: The maps of cross-correlation between the averaged temporal dynamics of the cortex (ROI 1 in Fig. 5) and the temporal dynamics of each voxel in the image, corresponding to Fig. 5.** The time lags represent different lags calculated from the cross-correlation function. ROI 3 represents the enlarged ROI in Fig. 5, and ROI 4 delineates the ventricle region. ROI: region-of-interest. The scale bar represents 1 mm.

### Supplementary Notes

#### Supplementary Note 1: Light refraction and focal distance calculation for the fUSPA imaging device

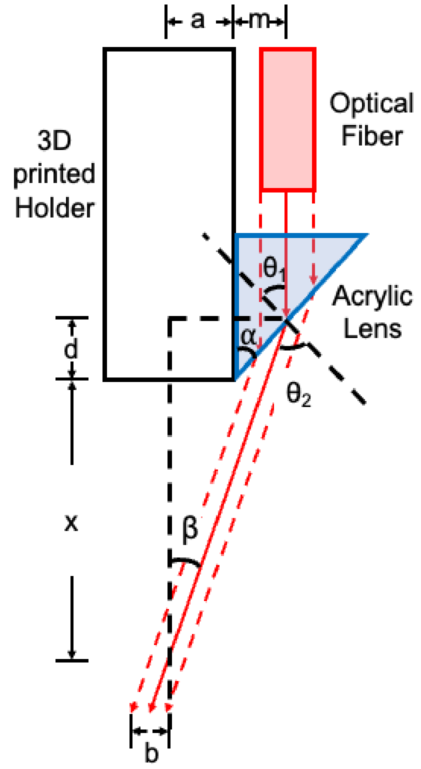


**Fig. S14: The schematic for the laser light refraction through the custom designed acrylic lens and associated focal distance calculations for the fUSPA imaging device.**

As shown in Fig. S1, two acrylic lenses attached to either side of the ultrasound probe, allowed the optical beam output from the fiber optic bundles to refract and co-align with the ultrasound detection zone (shown as the center black dashed line in above Fig. S14). This supplementary note calculates the half maximum width of the focal area ($b$) and the corresponding focal distance away from the transducer surface ($x$), based on the designed acrylic lenses with the slant angle a = 50°.

Assuming the optical beam output from the fiber normally incidents on the acrylic lenses and the optical profile is characterized in the air medium, according to Snell's law:

$$n_1 sin\theta_1 = n_2 sin\theta_2 \quad (1),$$

where $n_1$ is the refractive index for the beam input medium (acrylic lens) to be 1.4889 [1], and $n_2$ is the refractive index of beam output medium (air) to be 1. Substituting these numbers in, Equation (1) becomes:

$$sin\theta_2 = 1.4889 sin\theta_1 \quad (2).$$

As shown in Fig. S14, using the solid red line as light path and the reference, $\tan(\alpha)$ and $\tan(\beta)$ can be expressed as the following relationships:

$$\tan\alpha = \frac{m}{d} \quad (3),$$

$$\tan\beta = \frac{(a+m)}{x+d} \quad (4),$$

where m is the distance between the outer wall of the ultrasound probe holder and the input beam (red solid line). The optical fiber has a width of 2 mm and shielded inside a 0.5 mm thick stainless-steel casing. Therefore, the minimum $m$ is 1 mm and the maximum $m$ is 3 mm, respectively corresponding to the left and right red dashed light path as shown in Fig. S14. For the designed holder, dimension $a$ was measured to be 3.65 mm. Substituting the dimensions and combining Equation (3) and (4), $x$ can be expressed as:

$$x = \frac{3.65+m}{\tan\beta} - \frac{m}{\tan\alpha} \quad (5).$$

Based on the right triangles shown in Fig. S14, the relationships between $\alpha, \beta, \theta_1$, and $\theta_2$ can be written as:

$$\theta_1 = 90° - \alpha \quad (6),$$

and $\theta_2 = \beta + \theta_1$ (7).

Using Equations (2), (5), (6), and (7), and $\alpha$ = 50°, $x$ values are found to be 6.28 mm and 7.66 mm for $m$ values of 1 mm and 3 mm, respectively. Therefore, $x$ ranges from [6.28, 7.66] mm with the given fiber and ultrasound probe holder design.

The half maximum focal width ($b$) can then be calculated by the following equation based on similar triangles:

$$\frac{b}{a+m} = \frac{x_{m=3} - x_{m=1}}{x_{m=1} + d_{m=3}} \quad (8),$$

where $a$ is 3.65 mm, $m$ is at its maximum of 3 mm, and $x_{m=3}$ denotes the distance $x$ when $m = 3$ mm. Substituting the numbers into Equation (8), $b$ is calculated to be 1.05 mm. Therefore, given the lens angle of 50°, the maximum focal zone coverage width is $2b$ = 2.10 mm at x = 7.66 mm distance away from the transducer surface.

#### Supplementary Note 2: fUSPA system characterization and validation

We characterized and validated the multimodal imaging performance of integrated fUSPA system through multiple stages, including imaging of flat metal target, resolution wire targets, and flowing blood phantom. This supplementary note provides additional details regarding the characterization and validation results, in addition to those discussed in the Results and Online Methods sections of the main manuscript.

##### Pulse echo and photoacoustic A-line characterization

First, we conducted the US pulse-echo and photoacoustic signal characterizations to understand the sensitivity and element-to-element variation of the ultrasound probe. Accordingly, as shown in Fig. S3, the fUSPA system was used to image a thick stainless-steel flat target to characterize the US pulse-echo and PA A-lines. With the transducer surface placed parallel against the metal target (confirmed by Fig. S3a), the US pulse echo and PA A-lines were acquired from each element (the example A-lines for element 64 was shown in Fig. S3b). By calculating the peak-to-peak amplitudes of the A-lines of each element, we found an average amplitude of 1.33 ± 0.27 $x10^4$ (a.u.) for US pulse-echo and 1.49 ± 0.19 $x10^4$ (a.u.) for PA, respectively (Fig. S3c). Then, we further calculated the frequency spectrum corresponded to each A-line to understand the center frequency and bandwidth variances across different elements (the frequency response of the A-line from element 64 was displayed in Fig. S3d). As shown in Fig. S3e, we found an average center frequency of 15.20 ± 0.56 MHz (min: 13.65 MHz, max: 16.45 MHz) for US pulse-echo and 15.81 ± 0.74 MHz (min: 13.68 MHz, max: 17.45 MHz) for PA A-lines. The corresponding -6 dB bandwidths were found to be 40.31% ± 6.71 % (min: 27.61%, max: 58.62%) for US and 78.72% ± 6.97% (min: 62.12%, max: 97.18%) for PA, respectively (Fig. S3f). The -6 dB bandwidth from PA A-lines were roughly doubled than the US pulse-echo A-lines, which matched well with the difference between the one-way frequency response and the two-way frequency response, confirming the proper acquisition and sampling of the data.

Overall, the small variances in the US results demonstrated that the probe exhibited consistent acoustic performance across each element. The low variance PA results showed similar center frequencies as US but with doubled bandwidth, demonstrating that the fUSPA system produced uniform optical fluence that led to stable PA signal generation. These results indicated that the customized probe and its holder were suitable for high frequency USPA imaging applications with co-aligned light delivery and consistent acoustic transmit/receive performance.

##### Resolution and field-of-view characterization

Next, we conducted US and PA imaging characterization using the micro-wire based phantom to understand the resolution and the field-of-view (FOV) of the fUSPA system (Fig. S4, see Methods for details of the phantom). The resolution phantom is schematically displayed in Fig. S4a with three microwire targets placed at different depth inside a tissue mimicking phantom consisted of agar, silica, and intralipid. US contrast of the wire targets are generated from acoustic impedance mismatch with respect to background, whereas PA contrast was generated from the light absorption of the metal. By calculating the full-width-half-maxima (FWHM) of the point spread function (PSF) acquired through the second wire (W2) from both US (Fig. S4b) and PA (Fig. S4c) images separately, the lateral resolutions of the fUSPA system were characterized to be 189.9 μm (Fig. S4d) and 186.9 μm (Fig. S4f), respectively for US and PA. Similarly, the axial resolutions were measured to be 146.7 μm (Fig. S4e) and 193.7 μm (Fig. S4g), respectively for US and PA. PA imaging of micro wire targets distributed in the homogenous optical scattering medium showed a field-of-view (FOV) of 19 mm wide × 24 mm depth, without significant reduction in resolution or intensity.

**Flowing blood and static ICG tube imaging**

Afterwards, we validated the interleaved fUSPA system for its capabilities to image blood flow using a tube phantom (see Methods for details of the phantom). The phantom consisted of two 2.08 mm outer diameter silicone tubes with one flowing defibrinated bovine blood (Lampire biological laboratories, Pipersville, PA, USA) on the left and the other one with static indocyanine green (ICG) tube (1290 μM) on the right (Fig. S5a). While B-mode US was only sensitive to structural information and could not distinguish between the two tubes (Fig. S5b), UFUS based CBV mapping, which is highly sensitive to flowing particles, detected a strong signal from the flowing blood tube but no signal from the static ICG tube (right) (Fig. S5c). We also examined the PA signal, which is not sensitive to flow information but rather the optical absorption. We observed a stronger signal for the ICG tube comparing to the blood tube with the given wavelengths – 750 nm (Fig. S5d), 800 nm (Fig. S5e) and 850 nm (Fig. S5f). Moreover, clear differences were found in the intensity change between the flowing blood tube and the 1290 μM ICG tube across multispectral PA images, where the 1290 μM ICG tube intensity decreased with increasing wavelengths, while the blood tube PA intensity increased with increasing wavelengths. These results aligned well with the theoretical optical absorption curves of two molecules. To obtain the molecular content of the tubes, we used linear spectral unmixing to separate the oxygenated hemoglobin (HbO), de-oxygenated hemoglobin (HbD), and ICG contents. These results showed that the right tube had a strong unmixed ICG content (Fig. S5g), while the left tube had a high HbO content (Figs. S5h). We calculated the oxygen saturation (SO2) using the standard formula HbO/(HbD+HbO) x 100%, as displayed in Fig. S5i, and observed a clear distinction between the blood tube and the ICG tube. In addition, by supplying a gas mixture of $O_2$ and $CO_2$ at different concentrations to the flowing blood to mimic different oxygen saturation states for the blood, we validated the SO2 imaging capabilities of the fUSPA system. Using linear spectral unmixing, we observed a decline in the unmixed oxygen saturation as the oxygen concentration in the gas mixture reduced (Figs. S5j-m). Additionally, we examined the normalized PA intensity across the three wavelengths for ROI 1 (left side blood tube) in the case of 100% $O_2$ (blue solid line in Fig. S5n) and 100% $CO_2$ (blue dashed line in Fig. S5n), as well as for ROI 2 (right side 1290 μM ICG tube). The difference between 100% $O_2$ and 100% $CO_2$ was consistent with the assumption that the HbD content increased significantly in 100% $CO_2$ and dominated the PA intensity change. Additionally, the sharp drop of the intensity in ROI 2 followed the theoretical ICG in water optical absorption map at a concentration of 1290 μM. Overall, our findings suggested that the fUSPA system could provide quantifiable imaging of flow information from UFUS, while PA could extract the exogenous (e.g., ICG) and endogenous (e.g., HbO and HbD) molecular information with relative concentration changes.

### Supplementary Movies

#### Movie S1

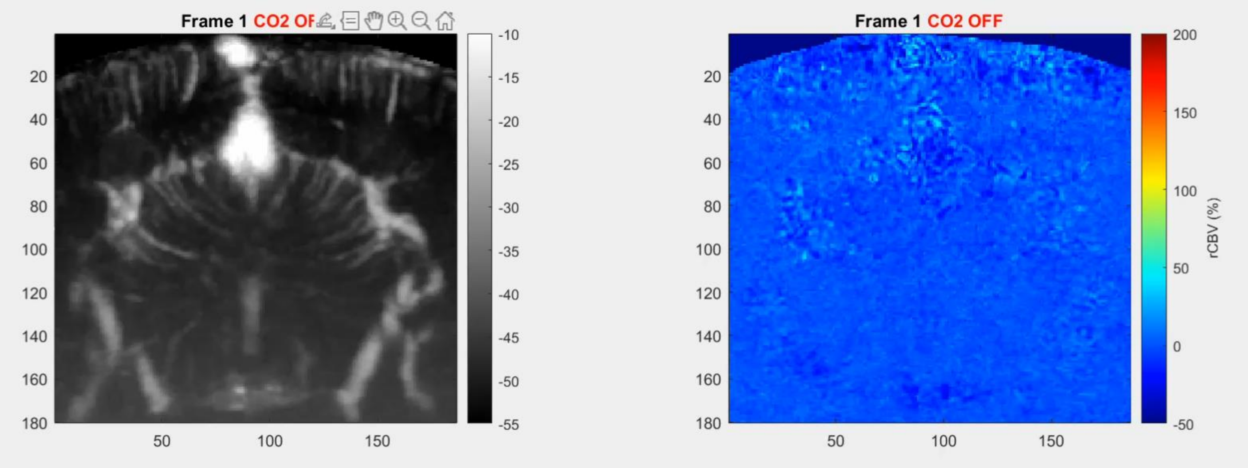


**Movie S1: Ultrafast ultrasound based cerebral blood volume (CBV) imaging of rat brain and relative CBV change map during hypercapnia stimulation, corresponding to Fig. 4.**

**Movie S2**

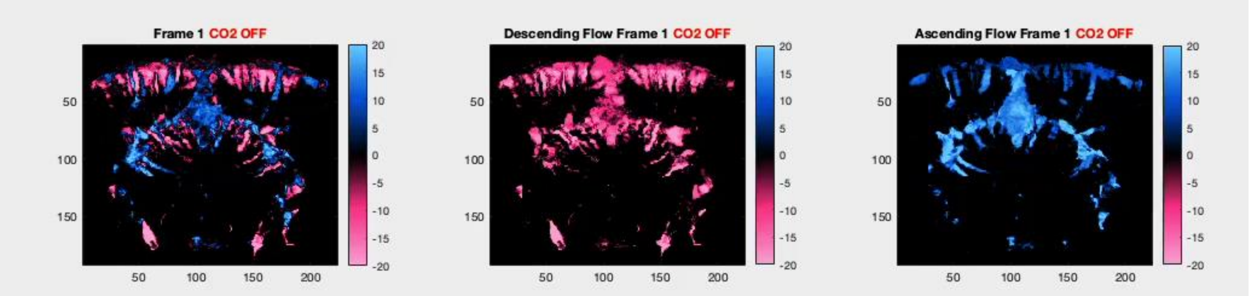


**Movie S2: Ultrafast ultrasound based descending and ascending cerebral blood flow maps in brain during hypercapnia stimulation, corresponding to Fig. 4.**

**Movie S3**

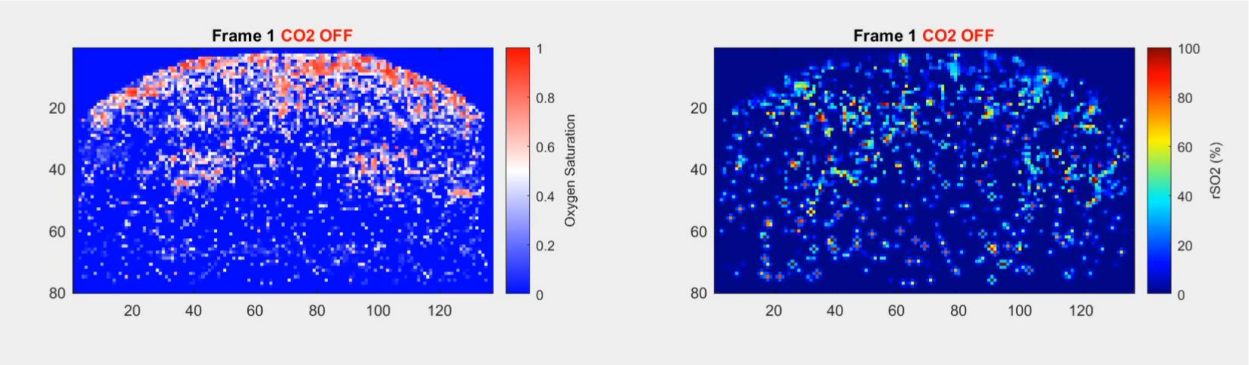


**Movie S3: Multispectral photoacoustic derived oxygen saturation and relative oxygen saturation change map during hypercapnia stimulation, corresponding to Fig. 4.**
